# Desert Tortoises in the Genomic Age: *Population Genetics and the Landscape*

**DOI:** 10.1101/195743

**Authors:** H. Bradley Shaffer, Evan McCartney-Melstad, Peter Ralph, Gideon Bradburd, Erik Lundgren, Jannet Vu, Bridgette Hagerty, Fran Sandmeier, Chava Weitzman, Richard Tracy

## Abstract

The California Department of Fish and Wildlife (CDFW) provided research funds to study the conservation genomics and landscape genomics of the Mojave desert tortoise, *Gopherus agassizii*, in response to the Desert Renewable Energy Conservation Plan (DRECP). To do this, we consolidated tissue samples of the desert tortoise from across the species range within California and southern Nevada, generated a DNA dataset consisting of full genomes of 270 tortoises, and analyzed the way in which the environment of the desert tortoise has determined modern patterns of relatedness and genetic diversity across the landscape. Here we present the implications of these results for the conservation and landscape genomics of the desert tortoise. Our work strongly indicates that several well-defined genetic groups exist within the species, including a primary north-south genetic discontinuity at the Ivanpah Valley and another separating western from eastern Mojave samples. We also use existing desert tortoise habitat modeling data with a novel extension of genetic "resistance distance" using geographic maps of continuous space to predict the relative impacts of five proposed development alternatives within the DRECP and rank them with respect to their likely impacts on desert tortoise gene flow and connectivity in the Mojave. Finally, we analyzed the impacts of each of the 214 distinct proposed development area “chunks,” derived from the proposed development polygons, and ranked each chunk in terms of its range-wide impacts on desert tortoise gene flow.

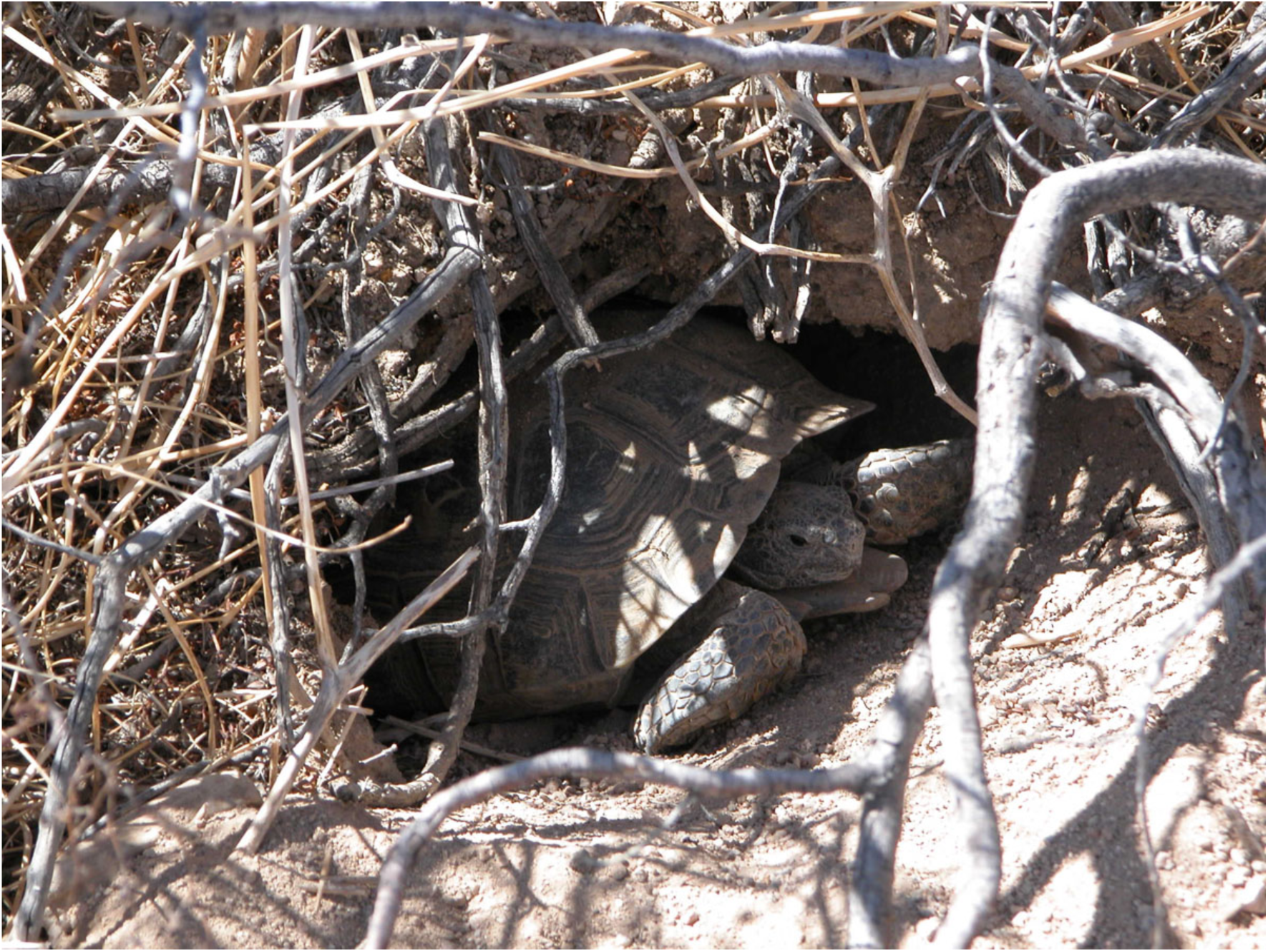

**Preface:** *Context:* The following document is a report that was submitted to the California Department of Fish and Wildlife, describing a series of analyses to help understand the impacts of several alternative spatial configurations of renewable energy development on gene flow of the federally threatened Mojave desert tortoise. These development alternatives were the centerpiece of the Desert Renewable Energy Conservation Plan (DRECP), a landscape-level land use planning initiative undertaken by the Bureau of Land Management (BLM), U.S. Fish and Wildlife Service (USFWS), California Energy Commission (CEC), and the California Department of Fish and Wildlife (CDFW). We were tasked by the California Department of Fish and Wildlife with providing a detailed analysis of these alternative plans on desert tortoise gene flow, and submitted the report for the public comment period for the initial implementation of the DRECP.

*Future Plans:* Current state of landscape-level planning for the Mojave desert tortoise
The five proposed land use configuration alternatives analyzed in the subsequent report include public and private lands spread across several counties in California. Shortly after the end of the DRECP’s public comment period, the government agencies that developed the DRECP announced that they would be splitting its implementation into two phases: one that deals with land use decisions on BLM-controlled lands and one that deals with non-BLM areas (Sahagun 2015).
Phase I of the DRECP was approved by the Bureau of Land Management on September 14, 2016 (U.S. Bureau of Land Management 2016). This phase includes land use planning decisions for BLM-administered lands. Specifically, 388,000 acres of public lands were designated as development focus areas (DFAs). In applications for leasing lands for renewable energy development, DFAs will not require the same degree of environmental evaluation prior to permitting, as they’ve already been evaluated in the context of the DRECP. The application process for renewable energy development within DFAs will be streamlined to encourage development in these areas. Phase I also designated a total of 6,527,000 acres for natural resource conservation. This includes California Desert National Conservation Lands, Areas of Critical Environmental Concern, and Wildlife Allocations. A further 2,691,000 acres were designated for recreation under Phase I. Phase II of the DRECP is currently under development in conjunction with county-level governments to extend this landscape-level planning beyond BLM-administered lands.

*Author Contributions:* This was a collaborative report. Evan McCartney-Melstad performed the simulations of the low-coverage full genome approach (see Figure 2); conducted all of the laboratory work to generate the genome sequences; performed all of the bioinformatic analyses to bring the raw sequence data to the various stages required for different analyses; wrote the software to quickly estimate pairwise genetic relationships between individuals using read count data in low coverage sequence data (www.github.com/atcg/cPWP); performed some of the population genetic analyses; and wrote and edited several sections of the report. Peter Ralph (in collaboration with Gideon Bradburd and Erik Lundgren) invented and implemented the random walk-based gene flow model that we used to estimate reductions in gene flow due to development, and also developed the theory behind the read-based pairwise pi and genetic covariance estimation used here, in addition to writing and editing several sections of the report. Gideon Bradburd also performed some of the population genetic analyses and wrote and edited several sections of the report. Jannet Vu collected and curated the spatial environmental data and generated the maps that are included in the report (Figures 10, A10-A13), and also wrote Appendices I and IV. Bridgette Hagerty, Fran Sandmeier, Chava Weitzman, and C. Richard Tracy contributed approximately 1,000 desert tortoise blood samples that they collected (at great effort), in addition to knowledge of tortoise ecology and conservation, as well as the results of previous microsatellite-based genetic analyses and editing of the report. H. Bradley Shaffer wrote and edited several sections of the report, and is listed as the lead author for his role in conceiving of and obtaining funding support for the project.

## Introduction

The Mojave desert tortoise (*Gopherus agassizii,* recently identified as taxonomically distinct from the Sonoran desert tortoise, *Gopherus morafkai,* see Murphy *et al.* 2011) is a widely distributed but declining resident of the Mojave Desert. Potential renewable energy development may negatively influence future population trajectories of this species, placing it into direct conflict with renewable energy projects. Anticipated increases in direct mortality, habitat loss, and especially habitat fragmentation from renewable energy development within the Desert Renewable Energy Conservation Plan (DRECP) Planning Area need to be considered in combination with other population stressors in the desert tortoise’s range. Listed under both the federal and California Endangered Species Acts, the Mojave desert tortoise is and will continue to be a significant driver of reserve design under the DRECP. Gaining the most precise knowledge possible about tortoise population health, genetic diversity, population substructure, and exchange of migrants in this widespread, unevenly distributed species will contribute to a comprehensive conservation strategy for this and other species covered by the plan. It will also inform the delineation of ecologically meaningful reserves in California’s deserts and facilitate siting of viable zones for renewable energy that best accomplish the goals of energy development while minimizing impacts on current and future tortoise population dynamics.

Recovery of rare and endangered species requires a series of interrelated steps and approaches. One pressing need is to apply the best available scientific techniques to understand how widespread species are genetically connected across landscapes, and therefore the extent to which seemingly discrete populations are genetically and demographically connected or isolated. Acquiring these data via direct field observations (e.g. mark-recapture, radio transmitters) is time-consuming, challenging, and expensive, especially for a cryptic species like the Mojave desert tortoise, which occurs at low density, is frequently in underground retreats, and lives for decades. Both direct and indirect (genetic) measures of gene flow and connectivity should be used to fully realize how organisms traverse landscapes, and therefore how to best preserve historical connections in the face of human modifications and the habitat fragmentation that often follows. A combination of direct and indirect genetic analyses often brings complementary information to management. For example, direct measures may indicate daily and seasonal activity and movement patterns, whereas indirect genetic measures often better indicate the extent to which dispersal results in successful reproduction and the long-distance, but often infrequent, movement of genes across landscapes.

Until recently, the principal paradigm used to study how genetic variation traverses landscapes has been one of Isolation by Distance (IBD; Wright 1943). One nearly ubiquitous field observation is that genetic differentiation between populations increases with the geographic distance between them (Jenkins *et al.* 2010) (“Everything is related to everything else, but near things are more related than distant things;” Tobler 1970). With the advent of cheaper DNA sequencing and the rise of Geographic Information Systems (GIS) data availability, the field of landscape genetics has developed to quantify other aspects of environmental heterogeneity that shape patterns of dispersal and genetic variation above and beyond that derived from geographic distance alone. Recent innovations have brought inferences from the burgeoning fields of landscape ecology and associated ecological niche models (Elith and Leathwick 2009; Forman 1995; Peterson *et al.* 1999) into the equation, adding a much-needed GIS component to landscape genetics analyses. Fortunately, some of the groundbreaking research in this area has been carried out in desert tortoises. We briefly summarize that research below, before describing our methods and findings.

## Current state of knowledge of desert tortoise landscape genetics

Andersen and colleagues (2000) modeled tortoise density, measured using field observations, as a function of 11 spatial GIS data layers, and found that “soil composition and parent materials can be important determinants of habitat suitability.” In 2009, Nussear *et al.* expanded on this work, compiling a dataset of over 15,000 desert tortoise (both *G. agassizii* and *G. morafkai*) occurrence observations. They modeled desert tortoise presence and absence over its entire range as a function of 16 spatial GIS layers and built a habitat suitability model for the species.

At the same time, a parallel research program was generating DNA sequence data for desert tortoises to learn how gene flow between desert tortoise populations may be structured by landscape characteristics. Edwards *et al.* (2004) reported on genetic variation at seven microsatellite loci for 170 Arizona (*G. morafkai*) tortoises, some of which were also tracked via radiotelemetry. Their research indicated that “long-distance movements result in the exchange of genetic material among adjacent populations,” but that “estimates of gene flow predate anthropogenic habitat fragmentation and should not be taken as evidence that natural immigration/emigration still occurs.” That is, because of the long generation time of tortoises, it may take many years for genetic population substructure to reflect reduced patterns of migration caused by anthropogenic disturbances. They further warned that long-distance migrants may be a critical component of connectivity and metapopulation dynamics in the species.

Hagerty *et al.* (2011) used the habitat suitability model of Nussear *et al.* (2009) to parameterize a resistance surface and a least cost path map (where resistance to migration in a patch of habitat (McRae 2006) is the inverse of its suitability from the model). Using 20 variable microsatellite loci sequenced in 744 tortoises, Hagerty *et al.* (2011) found support for both an effect of geographic distance and topographical barriers in structuring patterns of gene flow over the range of the desert tortoise. Their summary figure showing resistance across the range of the Mojave desert tortoise suggested that many low-cost paths, including ones in and out of the Ivanpah Valley, existed among their 25 population samples. Latch *et al.* (2011) employed similar sampling (859 tortoises, genotyped at 16 microsatellite loci), and found support for the hypothesis that both natural landscape features (slope), and anthropogenic features (roads) were limiting gene flow.

These studies offered exciting clues into the biology of the desert tortoise and the way in which the species interacts with its landscape. They also provided the earliest foundational insights that can be gleaned from landscape genetic data within the landscape and conservation genetics communities for desert tortoises. However, these studies were also limited in their ability to draw strong, spatially explicit conclusions by a lack of genetic resolution and the inability to precisely model anthropogenic impacts on population genetic movement probabilities. Recent advances in next generation sequencing (NGS) have enabled the relatively cheap generation of many orders of magnitude more DNA sequence data, which in turn has enabled the detection and quantification of vastly more subtle landscape effects on patterns of current and historical gene flow, and we fully embrace these advances in the current work.

In the current study, our goal is to move beyond the foundational work that has been accomplished on Mojave desert tortoise landscape genetics and beyond what is achievable with SNP-based NGS studies. By generating results based on full genome sequencing, we bring a greater level of precision to landscape genetic analyses than has previously been possible for the Mojave desert tortoise. Rather than generate genetic data for a few thousand variable single nucleotide polymorphisms (SNPs) as is now sometimes done in NGS studies, we sequenced the entire genome (an estimated two billion base pairs) of 270 geographically distributed tortoises. The resulting dataset is, to our knowledge, the most comprehensively geographically sampled dataset of whole genomes in a wild species. We generated this massive DNA dataset from a sample of desert tortoises that evenly covers their range, used cutting-edge spatial statistics tools, and applied these tools to both an expanded collection of high resolution GIS data rasters (explained in detail in Appendix I) and the habitat model of Nussear *et al.* (2009) to quantify how gene flow between tortoises has been affected by historical landscape features and how it will be affected by future anthropogenic changes. Below, we discuss these results and present specific, actionable, and data-supported recommendations for the conservation of the desert tortoise.

## Deciding among genomic approaches

Researchers in population genetics now have a choice of many different types of data and strategies for genetic data collection. Traditionally, virtually all work has used single or multigene Sanger sequencing, and traditional data types have included microsatellites, mitochondrial DNA (mtDNA) and nuclear DNA sequence analysis. While Sanger sequencing is still considered the gold standard with regard to per-nucleotide accuracy, the amount of data generated in Sanger sequencing is limited due to the cost (it is very expensive on a per-nucleotide basis), labor, and time that such analyses often take to complete. Much more recently, several new techniques that take advantage of massively parallel next generation sequencing (NGS) platforms have begun to replace traditional Sanger sequencing approaches. These new approaches almost invariably rely on Illumina NGS technologies, and sequence much larger fractions of the genome in a single, highly parallelized sequencing experiment. Restriction site associated DNA (RAD) sequencing and targeted sequence enrichment are two recent advances that seek to isolate, sequence and analyze a consistent subset of the genome from each individual, often yielding several thousand informative SNPs. Microsatellites and RADseq are very useful for population genetic studies and can generate small (most microsatellite analyses) to larger (RADseq) amounts of informative data. However, both suffer from problems with missing data, null alleles (particularly for microsatellites), and a general lack of information about the genomic distribution of informative markers. Targeted sequence enrichment is a technically more complex approach, and relies on having existing genomic resources that allow a researcher to identify (“target”) specific genomic regions to study. Target enrichment studies can also generate thousands of SNPs whose physical location and linkage relationships within that genome are known, but at relatively high cost in comparison to RADseq.

The newest, and most radical, approach, and the one that we employed in this study, uses whole-genome sequencing to characterize the entire genome of each individual. Recently, sequencing technology has progressed to the point where entire genomes can be sequenced, at low coverage, for prices accessible to wildlife researchers. Uncertainty associated with the genotype of any SNP at any particular position in the genome in low coverage, whole genome sequencing is much greater than for other approaches, because RADseq and target capture generally sequence each site many times. However, whole genome sequencing provides information for many orders of magnitude more sites than microsatellites, RADseq or target capture, and the statistical power gained appears to far outweigh the uncertainty from low-coverage, whole genome approaches.

## Methods

### Sampling

Desert tortoise blood samples were obtained in the field between 2004 and 2013, from two sources: samples collected by Bridgette Hagerty, Fran Sandmeier, and Chava Weitzman, working in Dick Tracy’s research group, and material stored at the Desert Tortoise Conservation Center (DTCC). We used existing georeferenced data to map all available tissues, and based on a visual assessment of those samples and a day-long discussion with Dr. Kristin Berry (USGS Western Ecological Research Center) on high-priority areas in need of analysis, we chose 270 blood samples for genomic analysis (see Figure 1). Blood samples had been stored in tubes by themselves or in RNAlater (Life Technologies) and kept frozen at -80C, or as dried drops on filter paper at room temperature. All blood samples were transferred to the Shaffer lab at UCLA for genetic analysis.

**Figure 1:**
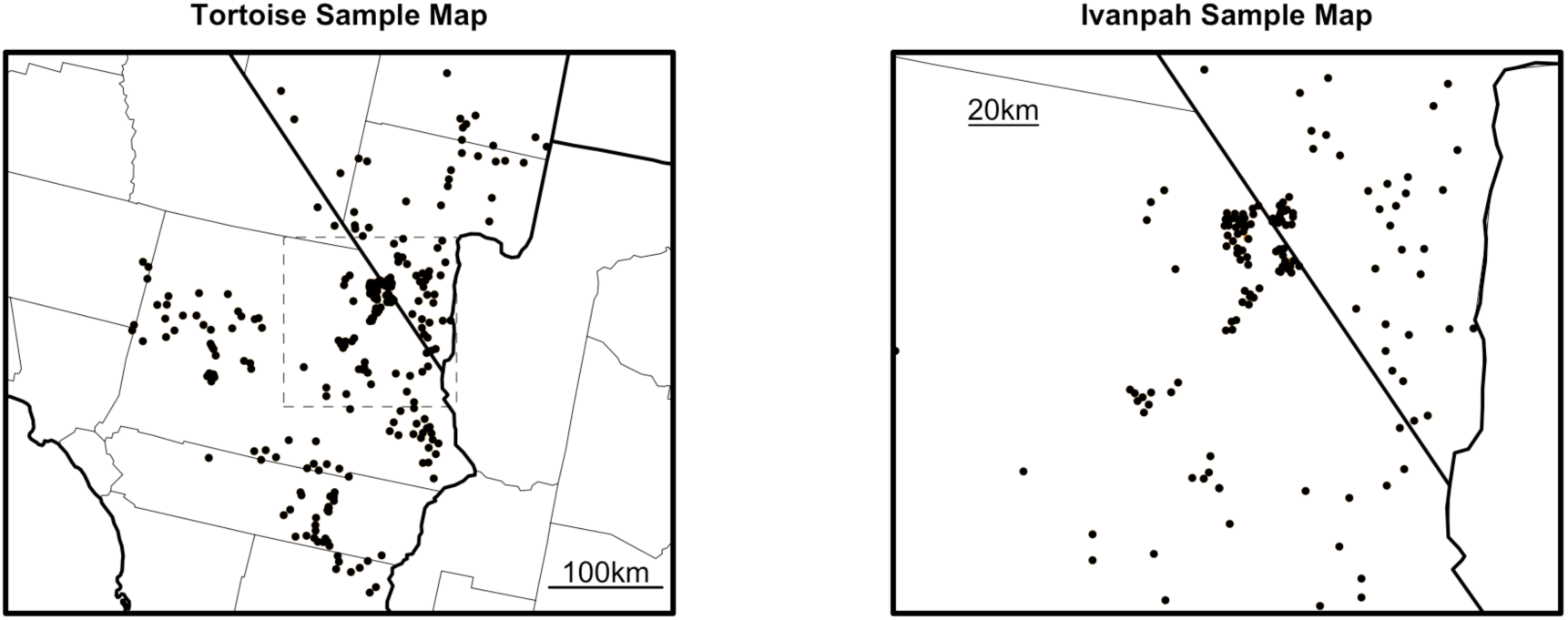
Sample map

### Laboratory Procedures

DNA was extracted using a salt extraction protocol (Sambrook *et al.* 2001), and quantified using the Quant-It dsDNA kit (Invitrogen). After diluting to 30ng/uL, DNA extractions were physically sheared to approximately 200-600bp using a BioRuptor (Diagenode) with an average of 7 cycles on the highest setting (30 seconds on, 90 seconds off). Sheared DNA was cleaned using a SeraPure bead mixture to remove chemical contaminants and to exclude short DNA fragments, which reduce sequencing efficiency. Illumina sequencing adapters were ligated to fragmented DNA using a standard library preparation kit (Kapa BioSystems). Each sample received one of 10 adapters with distinct 10 base pair indexes that allowed for samples from different tortoises to be combined, multiplexed in a single sequencing lane, and later computationally separated back into individual-tortoise data. All indexes had an edit distance of at least three to other adapters to allow for sequencing errors in the index reads (Faircloth and Glenn 2012). Importantly, the protocol did not include any PCR amplification, which is known to introduce and amplify biases in the resulting sequence data (Aird *et al.* 2011).

A total of 270 tortoises were submitted for sequencing at the Vincent J. Coates Genomics Sequencing Laboratory at UC Berkeley in three batches. Sample libraries were quantitated using qPCR and pooled, 10 samples per lane, for whole genome sequencing. All samples were sequenced in 100bp paired-end mode on an Illumina HiSeq 2000 or HiSeq 2500. Data for individual samples were de-multiplexed using Illumina’s CASAVA pipeline and downloaded in FASTQ format for analysis.

### Genomic Data

We developed a multi-step pipeline to remove low quality data, consistent with current best practices in processing genomic data. Particularly for low-coverage data (our data were approximately 1.5X coverage, which is very low), this is a critical step. First, reads that failed Illumina’s CASAVA filter were removed. Next, each sequencing read was checked for adapter contamination and base call quality degradation. Adapter contamination arises when the fragment being sequenced is shorter than the read length, resulting in bases being called at the 3’ end of fragments that are actually part of the synthetic sequencing adapters rather than the tortoise’s genome. Base call quality degradation occurs because the quality of base calls often deteriorates from 5’ to 3’ on Illumina sequencing reads (Fuller *et al.* 2009). To account for both of these factors, reads were processed using Trimmomatic 0.32 (Bolger e*t al.* 2014). Specifically, leading 5’ base pairs with a phred quality score (a standard metric of probability of accurate base-calling) below 5 were removed, and trailing 3’ base pairs with a phred quality score below 15 were removed. Then, a four base pair window was moved from 5’ to 3’ along each read, and the read was trimmed when the average phred base quality within the window dropped below 20. After this trimming, all reads less than 40bp in length were discarded.

Following sequence trimming, overlapping read pairs were merged using fastq-join from the ea-utils toolkit (Aronesty 2011). For short fragments, paired-end reads will overlap when the total length of the fragment being sequenced is less than two times the read length; merging these reads prevents artificially inflating sequencing coverage estimates where reads overlap, and results in improved mapping efficiency for these reads. Joined read pairs were then combined with singleton reads whose mate pairs were discarded in earlier quality control steps. This resulted in a set of paired reads and a set of singleton reads for each tortoise.

Paired reads and singleton reads were separately mapped to a draft of the Galapagos tortoise (*Chelonoidis nigra*) genome supplied by the laboratory of Dr. Adalgisa Caccone at Yale University. Ideally, reads would be mapped to a Mojave desert tortoise genome, since that would allow the maximum number of reads to be identified and physically arranged along the species’ genome. However, a high-quality genome for the Mojave desert tortoise did not exist at this stage of the project (although subsequent work will use an assembled desert tortoise genome (Tollis *et al.* 2017)). Mapping was done with bwa mem version 0.7.10-r998-dirty (Li 2013). Sequence alignment map (SAM) files output from bwa mem were converted to binary alignment map (BAM) and the paired and singleton alignment files for each tortoise were merged into a single alignment file for each tortoise using samtools version 1.0 (Li *et al.* 2009).

Merged BAM files were then cleaned to soft-clip alignments that extended past the end of reference contigs (CleanSam) and individual tortoise read group information was added (AddOrReplaceReadGroups) using picard 1.119 (http://broadinstittute.github.io/picard). Duplicates were then marked to identify levels of optical duplication (single molecule colonies on an Illumina flow cell that are mistakenly identified as multiple reads and can inflate coverage estimates). It was not necessary to mark or remove PCR duplicates because we utilized a PCR-free laboratory protocol, eliminating this potential source of error. Mapping rates were then calculated by counting the appropriate alignment flags using samtools flagstat (Li *et al.* 2009).

### Modeling whole-genome sequencing

Since low-coverage, whole genome sequencing is not commonly utilized, we first conducted a simulation study to compare the approach to RADseq. We simulated datasets from two hypothetical populations separated by a true genetic distance of Fst=0.001. In the first simulation, we sampled two thousand polymorphic loci (SNPs), at 20X coverage (typical for RAD sequencing or target capture). In the second, we simulated one million polymorphic loci, at 1X coverage (a low, and therefore conservative number of SNPs for whole-genome sequencing). The low, but real level of genetic divergence between these simulated populations was not recoverable using the first dataset, but was confidently inferred using the second (Figure 2). This result is consistent with a known guideline for population genetic analyses (Patterson and Reich 2006): genetic differentiation (measured by Fst) between two groups or individuals becomes detectable if it exceeds 1/sqrt(n*m), where m is the number of variable markers (∼1 million in our simulation) and n is the number of individuals. Based on these results, we were convinced that we should pursue a low-coverage, whole genome approach to best quantify Fst among individual tortoises.

**Figure 2.**
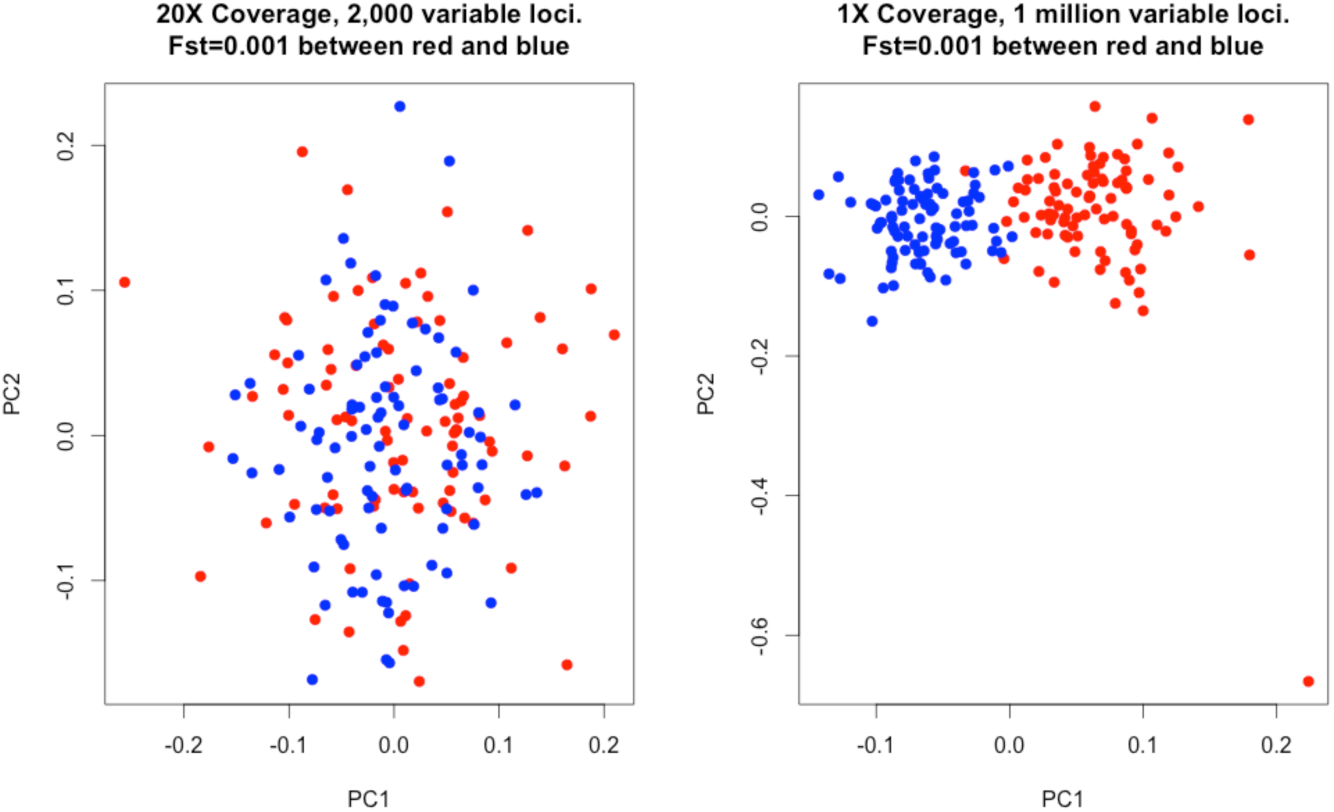
Comparison of two different sequencing approaches in their ability to differentiate very slightly differentiated populations (Fst=0.001)

### Inference of Genetic Relationships

In this study, the individual tortoise comprised our sampling unit. To determine the relationships between the sampled tortoises, we estimated two quantities for each pair of tortoises: pairwise sequence divergence, and genotype covariance. Pairwise sequence divergence is the average density of sites at which the two sequences differ, and is hence proportional to the average time back to the most recent common ancestor of the samples, averaged across the genome. Genotype covariance, on the other hand, decreases with average time back to the most recent common ancestor (Slatkin 1991), and is the basis for several widely used visualization methods.

To determine the set of variable sites used to estimate these quantities, we used *angsd (http://popgen.dk/wiki/index.php/ANGSD)*, an existing set of computational tools designed to incorporate uncertainty in genotype calls deriving from low-coverage sequencing data (Kim *et al.* 2011, Korneliussen *et al.* 2014, Li *et al.* 2010). This software provided us with reliable lists of polymorphic sites (SNPs), but unfortunately, we found that genotype posterior probabilities generated by *angsd* were influenced by both the variation in sequencing depth between samples and the distance of a given sample to the rest of our tortoise samples. (This is an expected byproduct of *angsd*’s statistical method to share power across samples.)

Both of these influences can lead to incorrect population inferences about tortoise biology, and therefore require correction. To do so, we developed a new method that instead uses raw read counts and is robust to differences in sequencing depth between individuals. To calculate divergence between a pair of tortoises in a way that is not influenced by sequencing depth, we estimated the probability for each base pair that two homologous reads drawn from the two tortoises are different at that base and averaged this across the genome, weighting by the read depths in those tortoises at that site (Appendix 2). Using this method, pairwise genetic divergence is not correlated with sequencing depth. For the following analyses we computed these pairwise sequence divergences using the full list of 52,740,529 sites determined to be polymorphic by *angsd* with a p-value less than 1e-6 (p<0.000001) and for which no tortoises had a read depth greater than 10 (to avoid overweighting repetitive regions, which can also skew summary statistics). These were then corrected to the proper genomic scale by multiplying by the density of polymorphic sites (an average of 2.98% in the 1.899 billion relevant bases of the reference genome).

The mean sequence divergence between two sequences provides an estimate of the mean time since they shared a most recent common ancestor, averaged across the sequence and multiplied by twice the average substitution rate (Hudson 2007). To make our results more interpretable, for the purposes of fitting models we converted sequence divergences to years by dividing by an estimate of twice the average nucleotide substitution rate. The substitution rate, 2.06×10^‒8^ percent divergence per year, was estimated by dividing the pairwise sequence divergence for a large set of genes between a tortoise (*Manouria emys*) and the painted turtle (*Chrysemys picta*) by a fossil-calibrated divergence time estimate between those two lineages. These two values were derived from a different large-scale turtle genomics project ongoing in the Shaffer lab. This estimate is probably not a completely accurate estimate of the true mean substitution rate for the desert tortoise, but is by far the best estimate currently available for turtles and tortoises from this related group of species, including the desert tortoise. It provides a reasonable, albeit approximate, idea of the time scales involved in our estimates.

We pursued multiple avenues of visualization and analysis of population structure. To investigate the geographic structure of genetic variation, we compared and plotted average pairwise sequence divergence against pairwise straight-line (Euclidean) distance between samples. In addition, we performed a principal component analysis (PCA) using the sample genetic covariance matrix, and obtained simple ‘geogenetic maps’ (inset of Figure 4) by plotting different PC axes against each other (Menozzi *et al.* 1978, Patterson and Reich 2006, Novembre *et al.* 2008). We also plotted the first several principal component scores for each tortoise onto elevational maps to visualize how tortoise genetic differentiation was distributed across their actual range (Figures A1 and A2).

### Isolation by Environment

It may well be that knowing simply where tortoises do not go (e.g. up steep mountains) suffices to describe gene flow across the species range. However, there are good reasons to suspect that other environmental factors have substantial effects. For instance, if overall habitat quality varies across the range so that there are “source” and “sink” populations, then we expect “source” populations to harbor more genetic diversity and to potentially serve as hubs connecting the “sink” populations. On the other hand, if offspring dispersal is biased such that young tortoises tend to end up in in habitats similar to their parents (beyond the correlation implied by localized dispersal), then gene flow between regions with different environmental variables will be reduced. This would imply that not only geographic distance but also ecological similarity predicts genetic differentiation, a pattern widely observed in nature (Sexton *et al.* 2013).

The current state-of-the-art method for making predictions of gene flow on continuous landscapes is to compute so-called *resistance distances* (McRae and Beier 2007). The nomenclature and formalism of this approach derives from a mathematical correspondence between electrical networks and certain quantities of reversible random walks. It turns out that if one equates the movement rates of a random walk between nodes in a network with the conductances of wires connecting those nodes, then the *effective resistance* between two points of the network (what one would measure using a volt meter) is equal to a biologically important parameter known as the *mean commute time* for the random walk, i.e. the mean time until a random walker, beginning at one of the points, first returns to its starting point after having visited the other point (Nash-Williams 1959). The results can depend on the discretization used and the resulting random walk model is metaphorical, not predictive. We used the fact that this correspondence, usually stated for discrete networks, carries over to continuous models, where the random walk is replaced by its continuous counterpart, a diffusion process whose movement rates depend on local properties of the inhomogeneous medium (in our case, the local landscape and its quality as tortoise habitat). This, combined with robust approximation of discrete random walks with continuous diffusions (Oblój 2004), allows us to bypass both drawbacks. The resulting resistance distance is then a powerful summary of gene flow across the landscape, since it integrates movement along all possible paths between the two locations.

The resistance distance has been shown to be a useful summary, but we would like to extract concrete predictions from it for effective management decision-making. Each generation since the most recent common ancestor provides an opportunity for mutations to occur that are inherited by only one of the sequences, and so mean sequence divergence provides an estimate of the mutation rate multiplied by the average time since the most recent common ancestor, across the genome (Hudson 2007). Focusing for a moment on a particular point on the genome, the time since the most recent common ancestor of two sequences for that part of the genome can be found by following the lineages back until they meet in their most recent common ancestor. Following a lineage backwards in this way can be seen as a random walk: the probability the lineage moves from location *x* to location *y* in a generation is the probability that a tortoise living at *x* has inherited the relevant bit of genome from a parent living at *y*. Intuitively, the motion of a lineage backwards in time looks like a random walk that is determined by the dispersal patterns of young tortoises, except that going backward in time, lineages are more likely to move towards better habitat, since more successful offspring are produced in such places. One important caveat is that it is known that under reasonable population models – in particular, those that show significant patterns of isolation by distance – the motions of two nearby lineages are not independent and therefore require a model that incorporates this non-independence (Barton, Depaulis, and Etheridge 2002). However, it is reasonable to assume that the motion of lineages is independent until the point that they are sufficiently close to each other in their path backwards to a common ancestor. Then, we can decompose the time to most recent common ancestor into two parts: the time until two lineages are close to each other, and the time from when they are close to each other until they find a common ancestor. This first part determines how sequence divergence decreases with distance, while the second part determines typical divergences between nearby individuals.

To relate this to resistance distance, we approximate the mean time until two lineages are close to each other by the average commute time. Specifically, we approximate the mean time until the lineages of tortoises at current locations *x* and *y* are within distance *d* of each other by one-half the sum of the mean time that a random walk begun at *x* takes to get within distance *d* of *y*, and the same quantity for *y* with respect to *x*. If the landscape is homogeneous then this approximation is exact, since the displacement between two independent walks is itself a walk that moves at twice the speed. On an inhomogeneous landscape it is a reasonable approximation, except in extreme circumstances (like very strong barriers to movement).

We provide the details and formal specification of this model of landscape resistance in Appendix 3. Briefly, we defined a random walk model whereby environmental rasters were each given two parameters that affect movement rates in the random walk: a stationary distribution and a relative jump rate to adjacent pixels. The application of a given set of stationary distribution and relative jump rate parameters (as well as a single overall scaling parameter) generates a resistance surface on the landscape over which commute times between tortoises can be measured. Since commute times are proportional to the coalescence times for pairs of tortoises, we can evaluate the model by testing how well random walk commute times over the generated resistance surface correlate with observed genetic distances. The optimal parameters for a set of landscape rasters are determined by minimizing the weighted mean square error for the set of tortoises used to fit the model.

### Landscape Resistance Models

We used the above procedure to fit a large number of landscape models, varying which landscape layers were used, which tortoises were used to fit the model, and which habitat mask was used. In all cases, we masked (that is, eliminated) regions to the east and south of the Colorado River, since they are now considered to be in the range of *Gopherus morafkai*. For reference, the tortoise habitat model of Nussear *et al.* (2009) fit a maxent model using 16 landscape variables, of which the most important were elevation (59.7%) and annual growth potential (AGP, 19.3%).

Each model fitting procedure produced a random walk model of tortoise lineage movement, which we evaluated in a common framework, measuring model fit to all tortoises using weighted median residuals. For an exact description, see Appendix 3 under Evaluating Landscape Resistance Models. We used median, rather than mean-squared, residuals to reduce the effect of statistical (and biological) outliers, and we weighted these so that the measure of goodness of fit assigns appropriate weights to each geographic area (unweighted would significantly upweight locations with more samples). Furthermore, we only use comparisons *within* each of the two major regions that we identified (north and south of the Ivanpah), because, as argued below in the results section on population structure, the relationship between the two regions has a deep-time historical component that is not likely to be a product of temporally homogeneous tortoise movement.

We evaluated a large number of possible landscape resistance models. Below is a quick summary of the procedure that led us to the best-fitting model.

First, we found that models fit using tortoises from both regions (loosely North and South) performed poorly: none could explain the two-cloud pattern seen in Figure 5. This is not surprising, because no available landscape layer accurately differentiates between those two regions. There is a confluence of not-insubstantial physical barriers around the break between the two regions (the mountains that define the Ivanpah Valley and the Colorado River), but the constriction in tortoise passage induced by these appears to not be sufficient to cause the genetic discontinuity that we detected. Furthermore, remaining tortoise population structure is seen to be much more significant in the north than in the south, and combining them into a single analysis confounds these differences. To deal with these differences we proceeded by fitting models using only comparisons between tortoises in the same group (north-north, or south-south comparisons). This is reasonable because we expect nearby comparisons to provide more information about local movement patterns than comparisons between tortoises on opposite sides of the range.

Next, we evaluated the effects of the choice of *habitat mask*, i.e. the region where movement was allowed to occur. We compared two choices: (a) the region for which the habitat model of Nussear *et al.* (2009) had habitat score above zero; and (b) the region below 2,000m in elevation. The first mask is strictly contained within the second; in both cases we also restricted to a reasonable bounding box (see the extent of the elevation layer in Figure 4). We found that the two different choices of mask gave indistinguishable goodness-of-fit values, and so proceeded with (a), the habitat mask based on Nussear *et al.* (2009), as this represents the best available biological prior knowledge of areas where tortoises are likely to avoid or perish, and which therefore should be excluded from our model.

Finally, we examined the impact of including different habitat layers in the model. We explored a wide variety of layers, but ultimately the best-fitting models all included only transformations of the habitat quality derived from Nussear *et al.* (2009). Therefore, we chose as our current best-fitting model the one providing the best goodness-of-fit using only transformations of the Nussear *et al.* (2009) habitat quality. (As discussed below, other models, including those with longitude, gave very similar results.) We favor this both because of its relative statistical simplicity and because it keeps our landscape model closely linked to the best available habitat model as derived by the desert tortoise biological community.

### Evaluation of Alternatives

We then used the best-fitting model to evaluate how development of particular areas under the DRECP would affect gene flow between different areas of the tortoise range. To do this, we evaluated changes in gene flow between each pair of a large set of reference points spread uniformly across the range predicted by Nussear *et al*. (2009), and we then used these to quantify both the overall reduction in gene flow and the areas that would be most affected (more details below). Some analyses considered “chunks” of proposed development areas within each of the five Alternatives separately. The process by which we generated these proposed development chunks is described in Appendix 4. In modeling how development on these “chunks” would affect gene flow, we assume that they represent zones of inaccessible habitat for tortoises, in the same way that areas outside the range boundary are modeled as inaccessible. Under our modeling strategy, a tortoise that wandered into a chunk boundary would reflect off of that boundary, much as it would if the development were surrounded by an impenetrable fence. Other modeling strategies are possible, and reasonable ones might include a pure mortality scenario (where tortoises have free access to development chunks, but die upon entry) or a semipermeable boundary (where some fraction of tortoises can cross the chunk). Given the uncertainty in exactly how development might occur in each chunk, we feel that our approach is a reasonable starting point, since it has minimal effects on demography (tortoises do not die when they reach a chunk boundary) but reasonable effects on gene flow (tortoises presumably cannot cross a large solar installation).

To quantify gene flow, we used the mean commute time to a 15-km circle (or neighborhood), since this is the same quantity used to fit the model. As discussed below, for a pair of points *x* and *y*, this is equal to one-half the sum of the mean time for a random walk from *x* to get within 15 km of *y*, and the mean time for a walk from *y* to get within 15 km of *x*. This can be concretely interpreted as the mean time since a tortoise at one location has inherited genetic material from a tortoise near the other location, along a particular lineage. Note that the neighborhood approach makes this measure independent of population density.

### Reference Locations

Our samples of tortoise tissue were not distributed uniformly across the range, and uneven sampling can have profound effects on some population genetic estimates and biological interpretations. To evaluate the effects of our sampling in an integrated way across the entire range, we chose uniformly spread reference locations as follows. First, we found the area with habitat quality of at least 0.3 in the Nussear *et al.* (2009) model, since those represented relatively high-quality tortoise habitat. Then, we sampled 10,000 points uniformly from across the enclosing rectangle, and discarded all but a maximal set of points that fell within the area of high habitat quality and had no two points within 10km of each other. This resulted in 202 points uniformly spread across the area of high-quality habitat. We additionally removed those points predicted by our model to be in isolated areas, defined as the minimal set of reference points such that after removing them, all remaining mean 15 km commute times were smaller than 3×10^6^ years (the maximum observed divergence between any pair of samples was slightly less than 1.5×10^6^, so a distance of 3×10^6^ would be equivalent to a separation of twice the width of the current range). The remaining points, shown on a map of habitat quality from Nussear *et al.* (2009), are shown in Figure 3.

**Figure 3.**
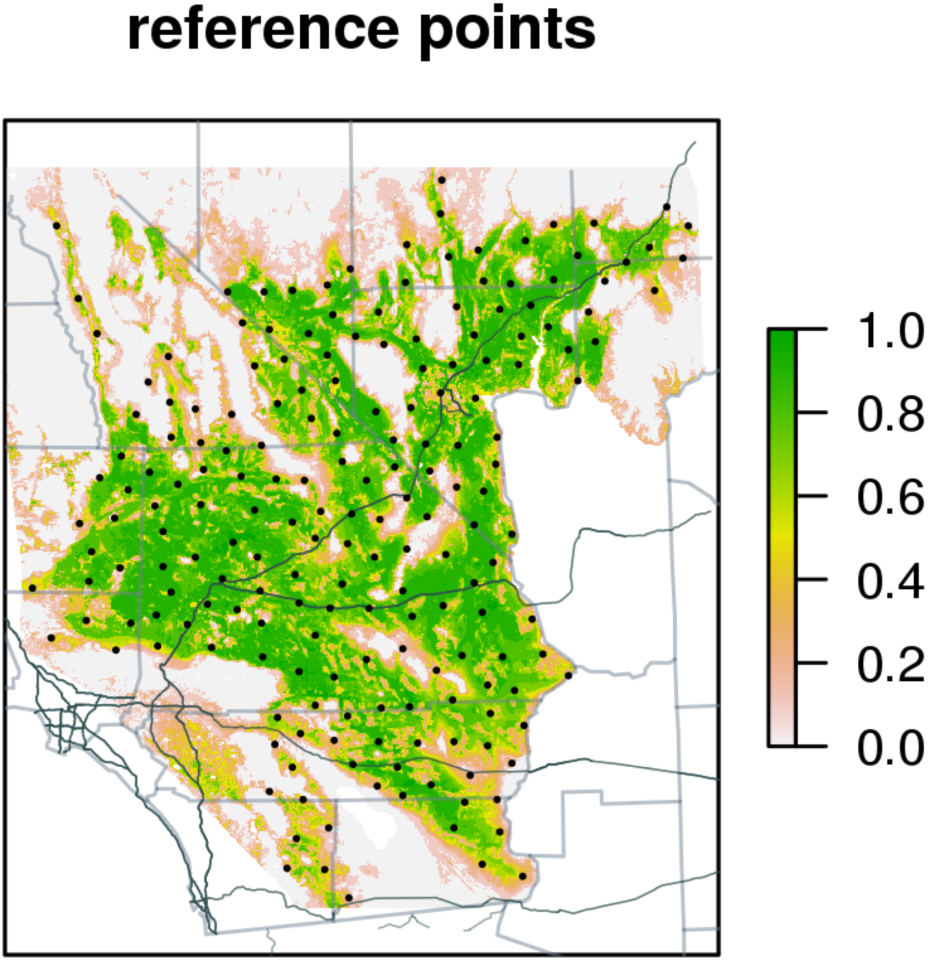
Reference points used to compute changes in gene flow across desert tortoise habitat.

### Measure of Isolation

The mean commute times described above allow us to quantify the effect that particular development scenarios will have on gene flow between any pair of locations in the range. However, this is not yet a measure of *isolation*. To quantify isolation, we need to summarize the total effects on each location across all of our sample points shown above. Consider, for instance, what would happen if a valley were to be blocked off to tortoises from the outside: mean commute times between the valley and the outside would drastically increase, but mean commute times within the valley would decrease, since tortoises within the valley can no longer take longer commutes outside the valley. Furthermore, and as we will see below occurs in practice, the act of removing a piece of habitat usually *reduces* commute times (or increases gene flow) between distant locations, because there are now fewer locations for transiting tortoises to visit. However, these commute times to very distant locations are not biologically relevant or important, at least on the time scale of human-mediated disturbances, simply because it takes thousands or millions of years for genes to commute to these distant locations, and that is not the scope of concern of our analyses. For these reasons, we say that a location becomes more isolated if mean commute time increases *to the bulk of the range*. We quantify this by identifying for each reference location the closest 40% of other reference locations, measured by commute time, and averaging the difference in commute time induced by removing a particular piece of habitat across those locations. This limits our summary statistics to a biologically reasonable region of space and time.

Concretely, suppose that 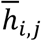 is the commute time between reference locations *i* and *j*, ordered by proximity to location *i*, so that 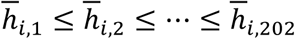. Then, since 202×0.4 ≈ 80, our measure of *isolation* of location *i* is

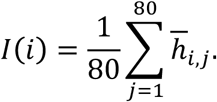

To interpolate values observed only on a subset of locations (e.g., the isolation values of the reference locations), we fit a thin plate spline model using the function *fastTps* in the *fields* package in R, which uses compactly supported kernels (with range 200km).

## Results

### Sampling

We amassed a collection of 270 tortoise tissue samples from throughout their range in the Mojave Desert (Figure 1). In our sampling, we attempted to balance an even spatial sampling of tortoises (increasing the probability of observing spatial structuring of genetic diversity partitioned by geography or environment) with a dense sampling of tortoises in regions of conservation importance (which also allows us to observe patterns of genetic differentiation on local spatial scales). The mean distance between a pair of tortoises was ∼141.8 km, with three-quarters of the distances less than 199.3 km, and ranging between 0 km and 464.8 km. The sampling was dense: the mean nearest-neighbor distance between tortoises was 6.4 km, three-quarters of the tortoises had another within 8.4 km, and only 10 did not have another tortoise within 20 km.

### Genetic Data

We obtained a total of 1.29 trillion base pairs of genomic sequence data from 28 paired-end 100bp Illumina HiSeq High Output lanes. Total bases sequenced per tortoise ranged from 1.71 billion bases to 13.91 billion bases, with a mean of 4.73 billion bases and standard deviation of 1.62 billion bases. Of this raw data, an average of 86.74% of reads passed Illumina’s CASAVA filter for each individual tortoise (standard deviation = 4.54%). After trimming low quality reads and merging overlapping read pairs as outlined in “Methods” above, the total number of bases going into the mapping stage ranged from 1.37 billion to 9.93 billion (average = 3.36 billion, sd = 1.12 billion).

### Mapping statistics

Mapping reads to the Galapagos tortoise genome was quite successful, with an average mapping rate of 95.67% (sd=0.67%). Using the ∼2.2 billion bp reference size of the Galapagos tortoise as a proxy for genome size, this yielded a mean sequencing coverage of 1.45X (sd=0.48, min=0.59X, max=4.28X).

### Population Structure

A geographically explicit way of looking at the relationship between genetic and geographic distance is to use PCA to summarize the major axes of genetic variation on a landscape. We show this in several ways. The positions of the samples on the first two principal components are shown in the inset of Figure 4. The most obvious pattern is the division of the samples into two large clusters by PC1, which corresponds to a fairly sharp division between tortoises to the north and south of the New York and Providence mountains (the eastern/southern border of the Ivanpah valley), with a few intermediate tortoises (coded as purple/pink) occurring in the Kelso area and the vicinity of Searchlight, NV. As the insert map of the Ivanpah region shows, these mountains form a strong barrier to tortoise dispersal; as a consequence PC1 accounts for about 12.2% of the total genetic variance in the data set. These two groupings also explain the two clouds of points that are evident in the overall IBD plot in Figure 5; genetic comparisons of pairs of tortoises between these two groups show a significantly higher divergence than comparisons of tortoises at comparable distances within each group.

**Figure 4.**
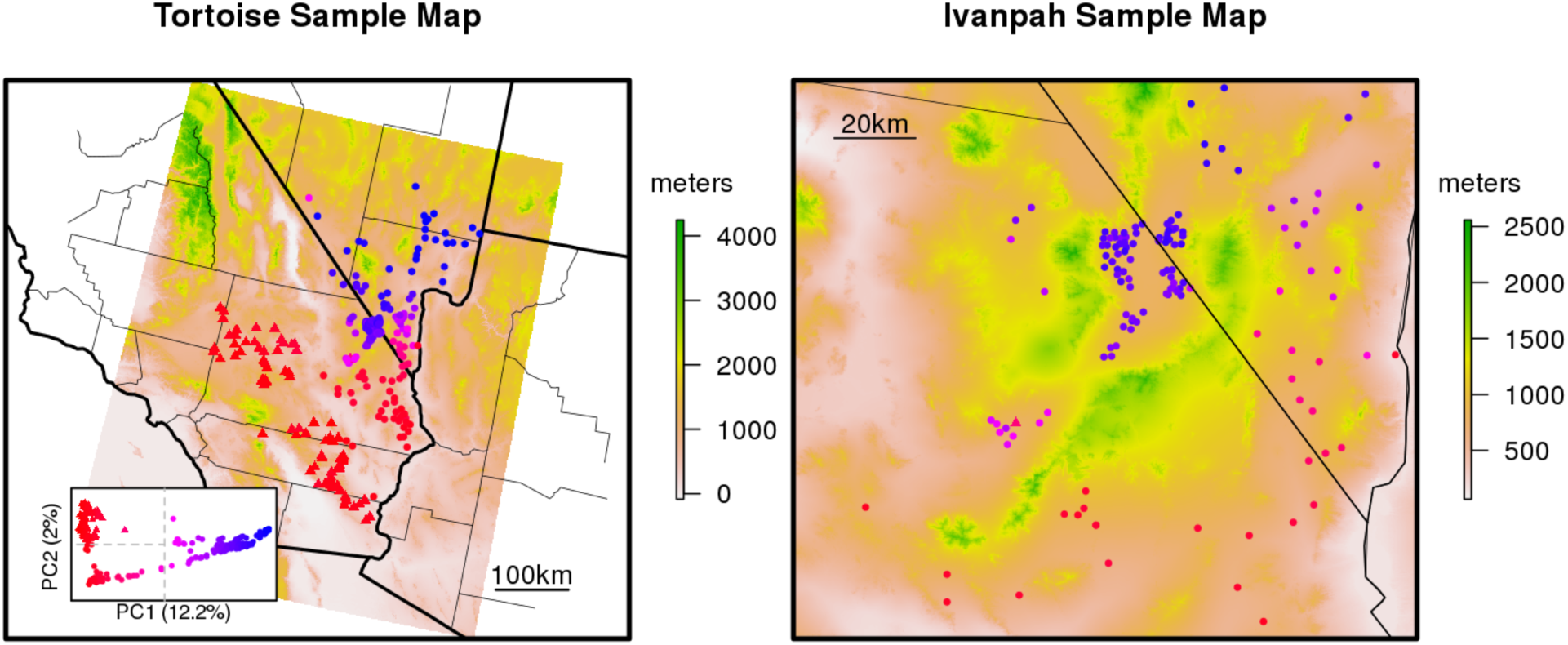
Tortoise sample map with samples colored continuously by their score on PC1. Additionally, samples on the left side of PC1 (mostly the red samples) were divided again into two sets based on PC2, with samples in the top half plotted with triangles and the other samples plotted in circles. The Ivanpah Sample Map is an expansion of this area that shows how genes move around the mountains in more detail. Background colors show elevation.

The first principal component is most striking, but others similarly reflect additional geographical subdivisions. PC2 further subdivides the southern tortoises roughly into eastern and western groups (red triangles vs. circles in Figure 4) on either side of the low-lying Cadiz valley lakebeds and accounts for about 2.0% of the total genetic variance. Subsequent principal components further subdivide the range, generally following geographical barriers such as mountains. These sub-groupings account for additional substructure seen in Figure 5, and represent geographic patterns of differentiation above and beyond that explained by distance alone. Overall, our emerging hypothesis is that relatedness between tortoises is well predicted by distance as traversed by tortoises on the landscape, with the exception of the north-south division, which may reflect the effects of past history.

### Isolation by Distance

The mean density of nucleotide differences between tortoises (“pairwise divergence”) in the sample is 5.4 differing sites per kilobase, and varies between 3.3 and 5.9 sites per kilobase. This measure of relatedness, when compared to geographic distances between tortoise pairs, demonstrates that tortoises sampled nearby each other are more closely related than ones sampled farther away (Figure 5) – the classic “isolation by distance” pattern (IBD; Wright 1943). Overall, pairwise divergence increases by about 0.0011 differences per kilobase for each additional kilometer of separation (Figure 5; p<10^‒16^). The “groups” referenced by Figure 5 are shown as discretely colored purple and blue dots in the left panel, and are determined by discretizing the samples by their scores on PC1 as shown in Figure 4.

**Figure 5.**
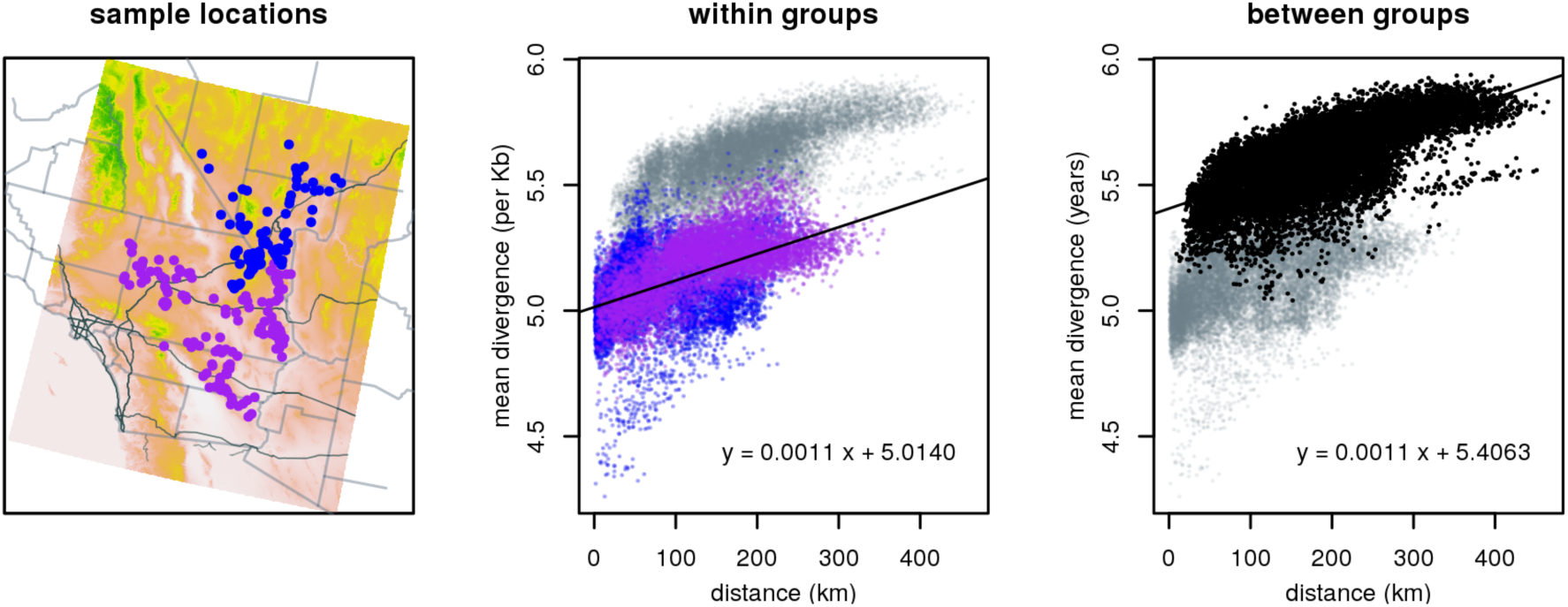
Showing the slope of isolation by distance both within each group and between the two groups. Groups are defined by their scores on PC1 and are shown in the left panel.

This positive correlation of genetic differentiation and geographic distance extends to the smallest spatial scales: within the Ivanpah valley, where the densest cluster of samples occurs, the relationship between pairwise divergence and geographic distance is likewise highly significant, showing an increase of 0.0024 differences per kilobase for every extra kilometer of separation (Figure 6; p < 10^‒16^).

**Figure 6.**
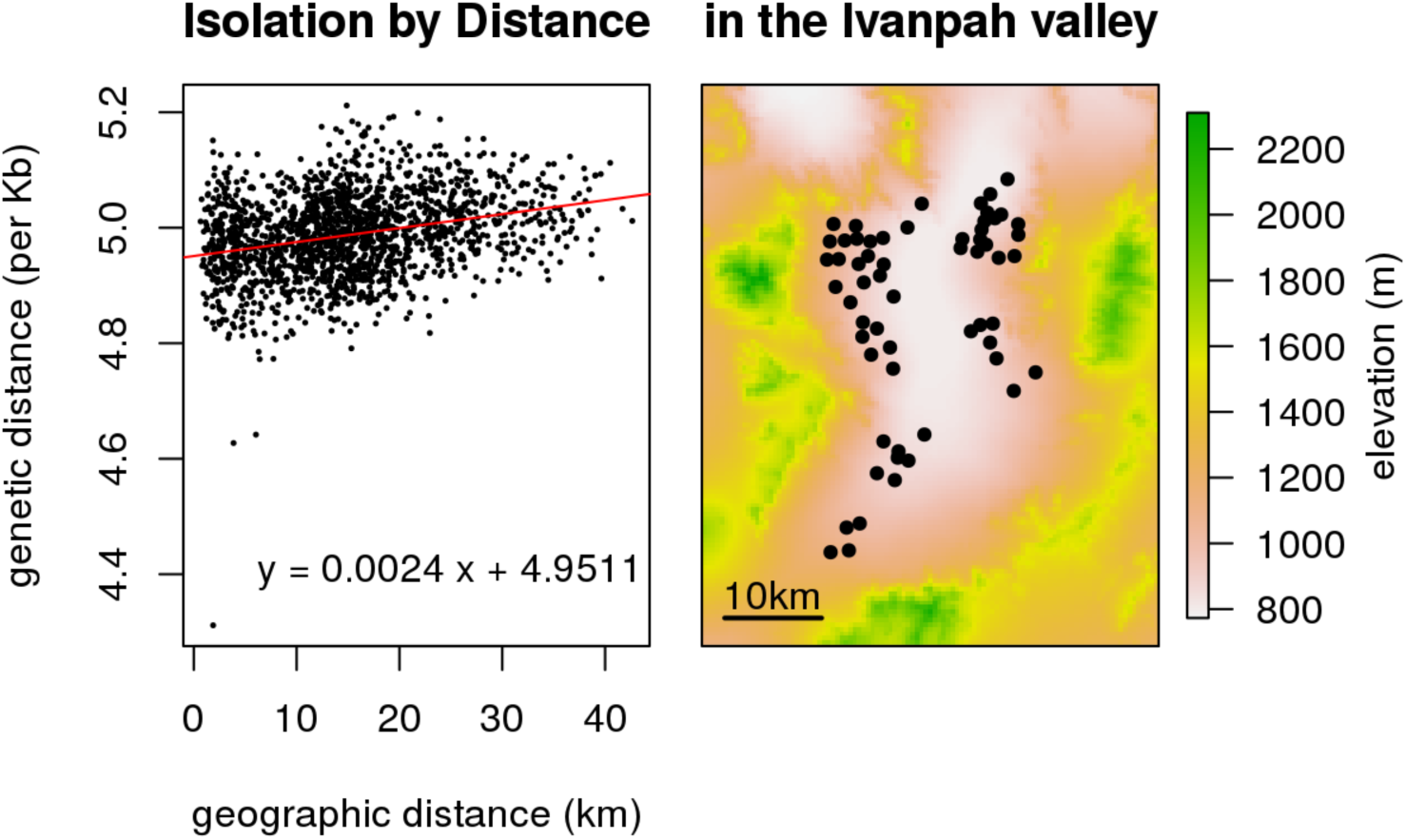
Pattern of isolation by distance solely within a small geographic range in the Ivanpah Valley. The map on the right shows the location of each tortoise sample.

### Model fit

Of the many models that we tested, the one with the best fit included the effects of only one layer: a binary layer that takes the value 1 if habitat quality (from Nussear *et al*. 2009) is greater than 0.3, and zero otherwise. (More extensive model fitting results are given below.) This model allows four rates of tortoise movement: on low quality habitat, on high quality habitat, from low to high quality habitat, and from high to low quality habitat. The weighted median residual value for this model was 2,133 years, while the same quantity for the linear regression of pairwise divergence against great-circle geographic distance was 18.7% greater. The difference measured by weighted mean squared error was even stronger: a 53.7% difference. This indicates that the landscape model incorporating a binary indicator of tortoise habitat quality as defined by Nussear *et al.* (2009) did a significantly better job of predicting genetic relationships between tortoises than did straight-line distance. Although additional, more complex models can and should be tested, we used this two-state model based on its simplicity, its biological realism, and its statistical performance.

### Effect on gene flow of removing habitat

As discussed in the *Methods*, we quantify the effects on gene flow of removing particular pieces of habitat through the changes in mean time for the random walk that models the time, in years or generations, that it takes for tortoise lineages to travel between each pair of points, averaged across reference locations. **These analyses provide the tools by which we can directly test the effects on tortoise connectivity of alternative habitat modification plans as outlined in the DRECP.**

### Effects on gene flow to single locations: examples

To show how this approach works, we consider how the removal of all of the habitat in the Preferred Alternative Plan would affect gene flow to a single location. In the left panel of Figure 7, the star located in the far western Mojave is the single location, and each map pixel is colored according to the mean time to reach the 15km circle surrounding the star in the map. Unsurprisingly, it takes longer to reach locations that are more distant from the star. On this map, the potential development areas of the DRECP Preferred Alternative are shown in grey, and the mapped mean times have been computed after blocking these areas to possible tortoise movement. The middle panel shows how this differs from the scenario where these development areas are not blocked: each area is colored according to the difference between the mean time to reach the starred area before and removing the development areas. As this map demonstrates, most parts of the range are around 40,000 years more distant (in red), although the nearby area that is also trapped between two potential development areas becomes slightly closer (in pale blue). The right panel shows the relative change: here, colors correspond to the difference (middle panel) divided by the mean time without the potential development areas removed. As might be expected, the relative effect is greatest near the star, since more distant areas are relatively less affected by a change near the star.

**Figure 7.**
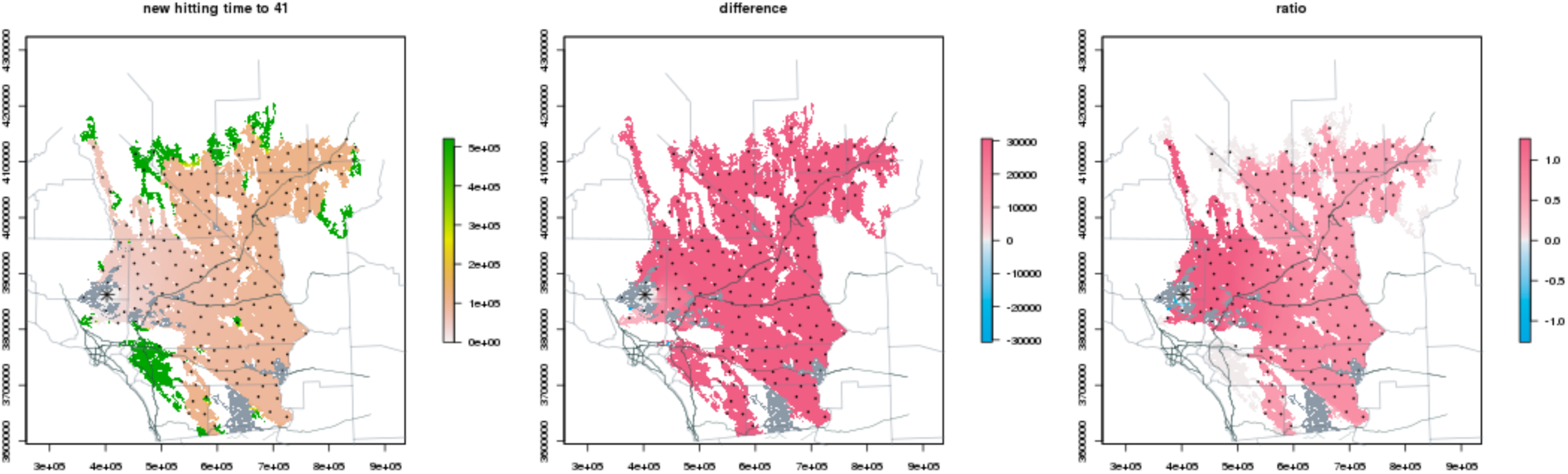
Showing the effects on hitting times to a single spot in the western Mojave as a result of removing the land in the preferred alternative. The dark green areas in the left panel were deemed inaccessible under this model.

Figure 8 shows the same set of analyses for a reference location near the center of the Mojave, again marked with a star. Here, we see that most of the Mojave actually becomes *closer* (blue in the center panel): this is because removing a portion of habitat means that there are fewer available locations for ancestors to live, and so all else being equal, two tortoises are expected to have ancestors living nearby to each other more recently. However, note that there is a small “shadow” of increased distance (red) just on the other side of a nearby potential development area, reflecting the reduced regional gene flow in this area that would be induced by blocking off this piece of habitat.

**Figure 8.**
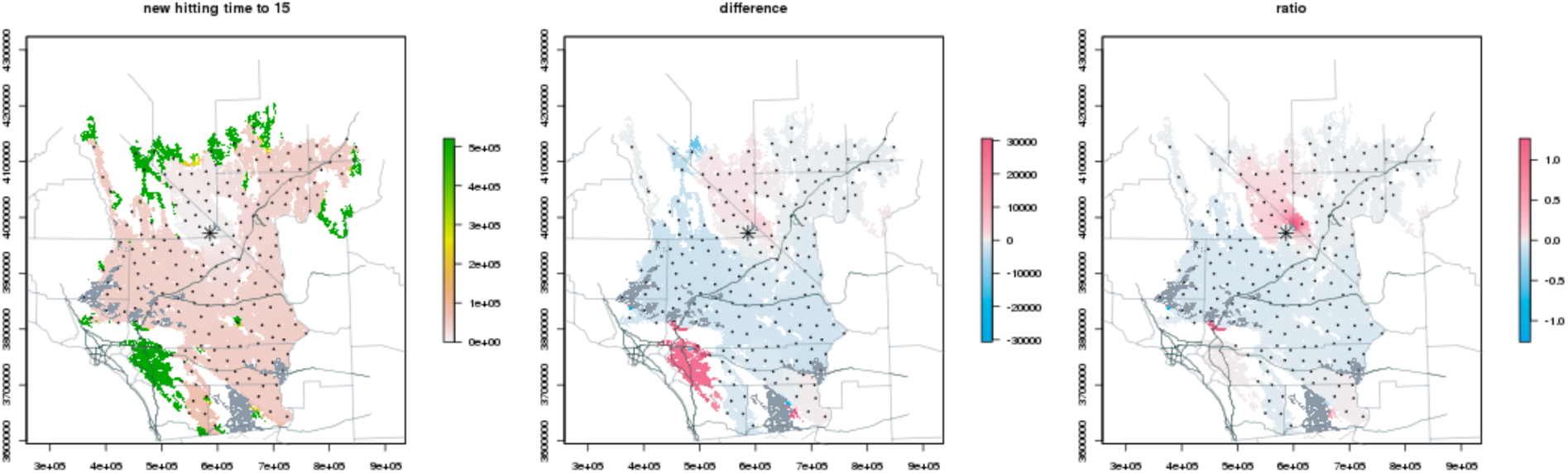
Showing the effects of hitting times to a single spot in the central Mojave as a result of removing the land in the preferred alternative.

### Combined effects on gene flow across the range

To summarize the effects on gene flow of blocking off particular regions of the habitat, we average the difference in gene flow with and without the possible barriers, across the nearest 40% of the other reference locations. We chose the closest 40% as a reasonable compromise between the entire range of high-quality habitat of the Mojave (which is too large to reasonably affect gene flow for a tortoise) and a region immediately surrounding an animal (which does not allow for the cascading effects across generations of blocking gene flow). We computed this measure of isolation for each reference location and interpolated it to the remainder of the map; this is shown on the left of Figure 9. We refer to this statistic as the *mean difference nearby*; it quantifies the mean difference in gene flow for the biologically relevant 40% of tortoise habitat with and without a subset of habitat removed. On the right we show the *relative difference nearby*, calculated as the mean (over the same 40% of locations) of the ratio of this difference in gene flow to the gene flow (commute time) in the original habitat without the barriers. These figures show the predicted impacts of removing the proposed development chunks of land in the DRECP Preferred Alternative.

**Figure 9.**
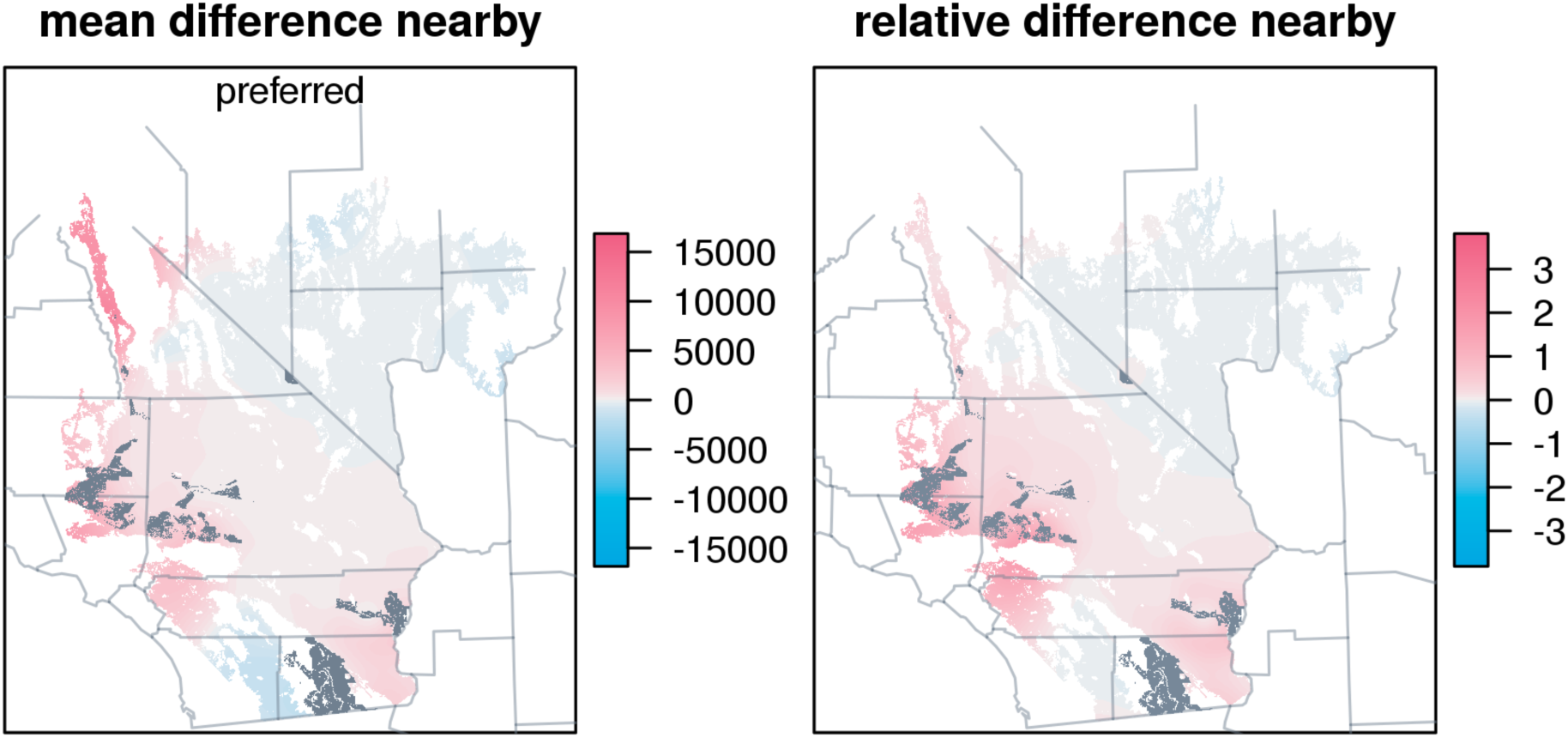
Mean and relative difference in commute times across the range as a result of removing the land in the preferred alternative.

In both maps, the darkest red areas are more distant from other, nearby portions of the range by 10,000-15,000 years. This is a very strong separation, because most parts of the range are separated by less than 10,000 years, as seen in the example commute time plots above.

### Comparing total effects of each alternative

We can now apply this same approach to each of the four Alternative plans in the DRECP, and compare them to the Preferred Alternative. We plot these in Figures A3-A6.

In order to be able to rank these alternative development plans, we show several summary statistics for each. For each alternative, areas are given in km2 and as a percentage of the total tortoise habitat. We calculated and tabulate each of the following:

- *habitat removed* is the total amount of area either in possible development areas or completely isolated from the rest of tortoise habitat under that alternative (this occurs if, for example, land is developed in a ring, with an undeveloped hole in the center of the development)
- *isolated* is the total area for which the gene flow to nearby areas has increased (regardless of the amount by which it has increased)
- *isolation* is the mean amount by which the commute time has increased to nearby areas across this area where it has increased; **it measures the intensity of decrease in gene flow**
- *strongly isolated* is the total area to which gene flow has strongly decreased; we define “strongly” as the mean commute time to nearby areas increasing by at least 1,500 years
- ***relative isolation* is the ratio of the amount by which commute time has increased to the commute time without blocking any areas, averaged over the set of nearby locations**

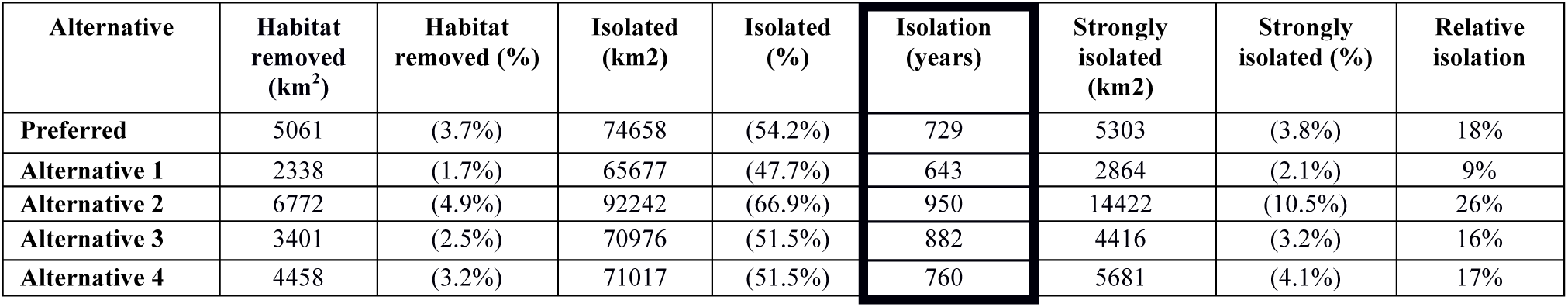

| Alternative | Habitat removed (km<sup>2</sup>) | Habitat removed (%) | Isolated (km<sup>2</sup>) | Isolated (%) | Isolation (years) | Strongly isolated (km<sup>2</sup>) | Strongly isolated (%) | Relative isolation |
| --- | --- | --- | --- | --- | --- | --- | --- | --- |
| Preferred | 5061 | (3.7%) | 74658 | (54.2%) | 729 | 5303 | (3.8%) | 18% |
| Alternative 1 | 2338 | (1.7%) | 65677 | (47.7%) | 643 | 2864 | (2.1%) | 9% |
| Alternative 2 | 6772 | (4.9%) | 92242 | (66.9%) | 950 | 14422 | (10.5%) | 26% |
| Alternative 3 | 3401 | (2.5%) | 70976 | (51.5%) | 882 | 4416 | (3.2%) | 16% |
| Alternative 4 | 4458 | (3.2%) | 71017 | (51.5%) | 760 | 5681 | (4.1%) | 17% |

Several points are worth noting in this table. First, as a single summary statistic of the effect of a development alternative, we highlight the Isolation (in years) column of the table, which summarizes the overall increase in isolation, or the decrease in gene flow, for each alterative. Second, every alternative increases the time for genes to traverse the landscape by many hundreds of years, approaching 1000 years for Alternative 2. These are large numbers representing very significant effects across dozens of tortoise generations. Third, Alternative 1 is clearly the least harmful (which makes sense given that it is the smallest acreage), and Alternative 2 is the most harmful. And fourth, the Preferred Alternative has a very substantial effect on tortoise connectivity (729 years).

**We also call attention to the Relative Isolation, particularly in comparison to the percentage of habitat removed.** In most cases, the effect in terms of Relative Isolation is about five times the percentage of habitat removed, reflecting the extremely strong, cascading effects that development has on tortoise movement. For example, for the Preferred Alternative, removing 3.7% of the tortoise habitat leads to an 18% increase in relative isolation.

### Effects of removing each chunk

Using the same analytical approach, we also consider the effects of removing each “chunk” of habitat for the Preferred Alternative. Figure 10 is a key showing where each chunk is found under the Preferred Alternative. The *mean isolation* is a reasonable measure of the total effect of removing a given chunk, but it must be interpreted with some caution. In particular, the absolute size of the *mean isolation* **only** measures this chunk, in isolation, without the effects of any other chunk that might be removed. There are many instances where the impacts of multiple chunks considered together have a much larger impact than the sum of the chunks individually, reflecting the synergistic negative effects that can result when multiple chunks are removed. We provide this table to show how individually chunks may have very different impacts; to understand the impacts of removing a set of chunks, **each potential combination of removal areas must be modeled and evaluated**, and we have not done that here.

We calculated and tabulate each of the following:

- *habitat removed* is the total amount of area either in possible development areas or completely isolated from the rest of tortoise habitat for this chunk
- *isolated* is the total area over which the gene flow to nearby areas has increased
- *mean isolation* is the mean amount by which the commute time has increased to nearby areas across this area where it has increased; it reflects the decrease in gene flow for each chunk in the analysis
- *max isolation* is the maximum amount by which the commute time has increased between any two nearby reference locations given the removal of this chunk

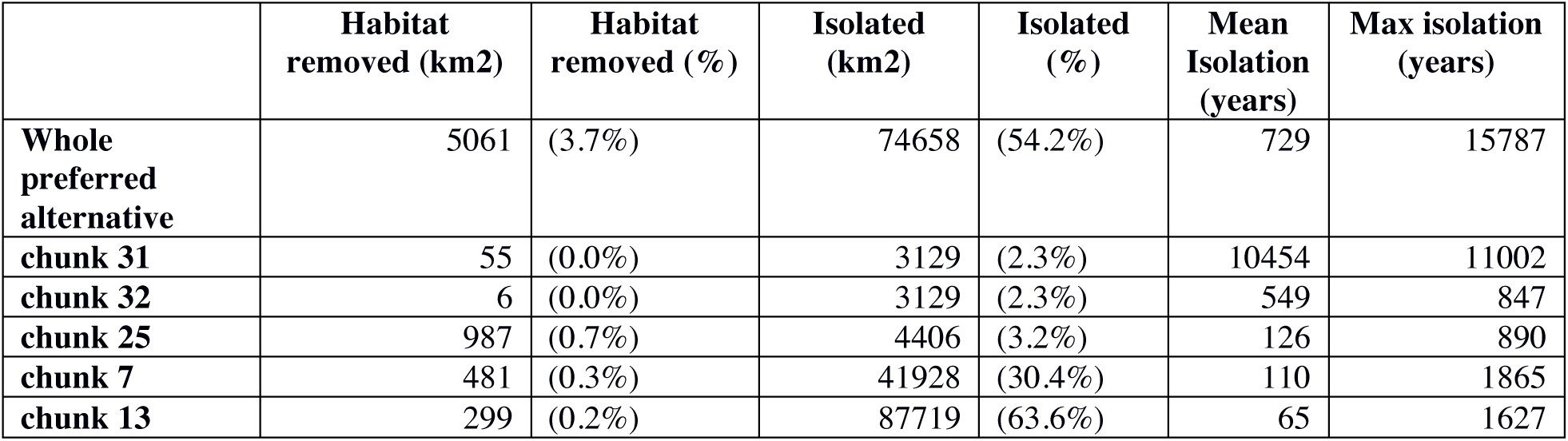

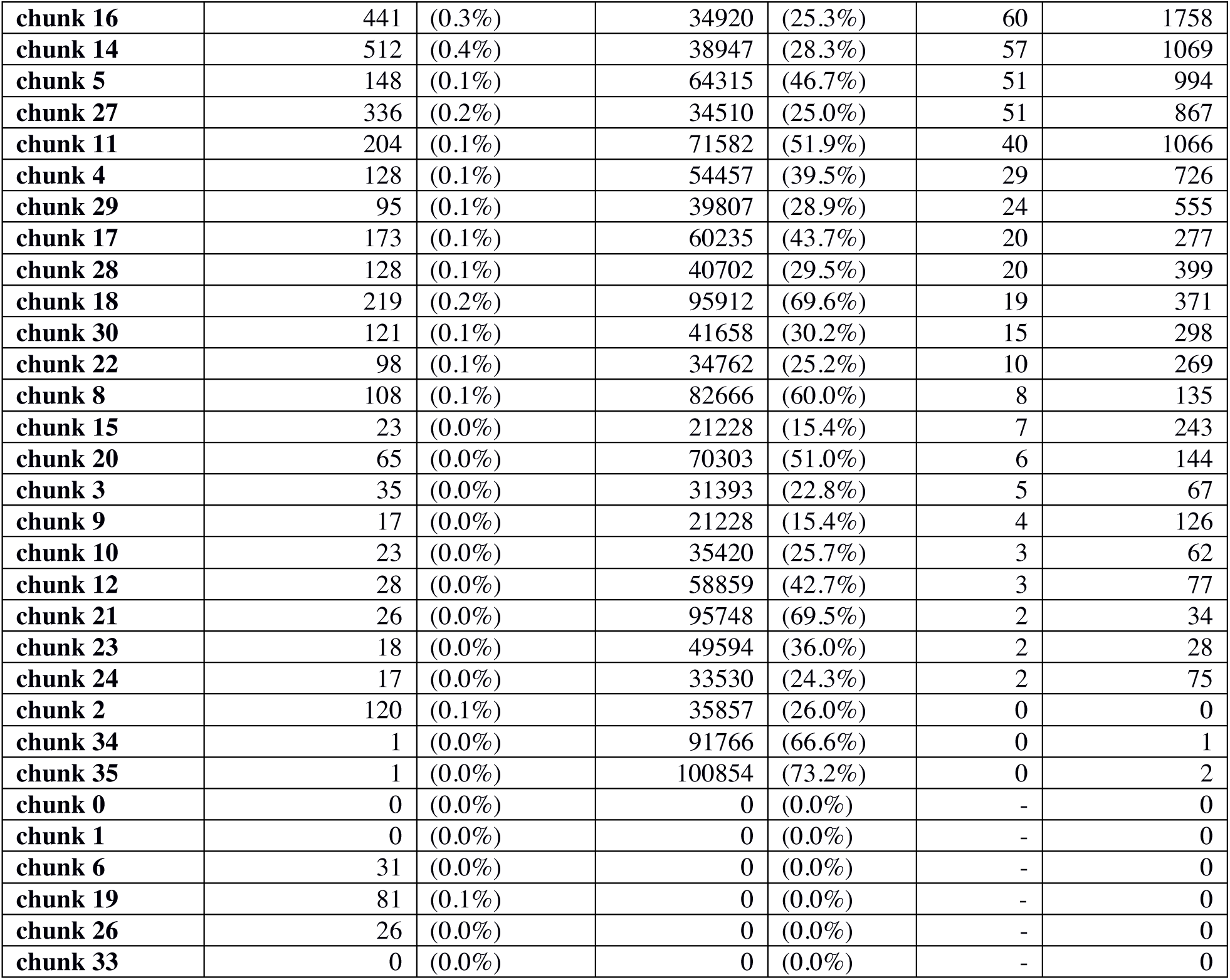

Here, the chunks are ordered by their Mean Isolation effect, from largest (most detrimental) to smallest (least detrimental). The primary point to take from this analysis is that both the amount of habitat removed and its spatial configuration are important determinants of the effect of a chunk, or project, on tortoise gene flow. For example, chunks 31 and 32 are both quite small in terms of area, but have extremely large effects on tortoise gene flow as reflected in their *mean isolation* effect size. This presumably reflects their position near the mouth of the Owens Valley, and their effective isolation of that entire piece of tortoise habitat. In contrast, chunk 25 is relatively large, but has a much smaller effect on *mean isolation* than chunks 31 or 32.

**Figure 10.**
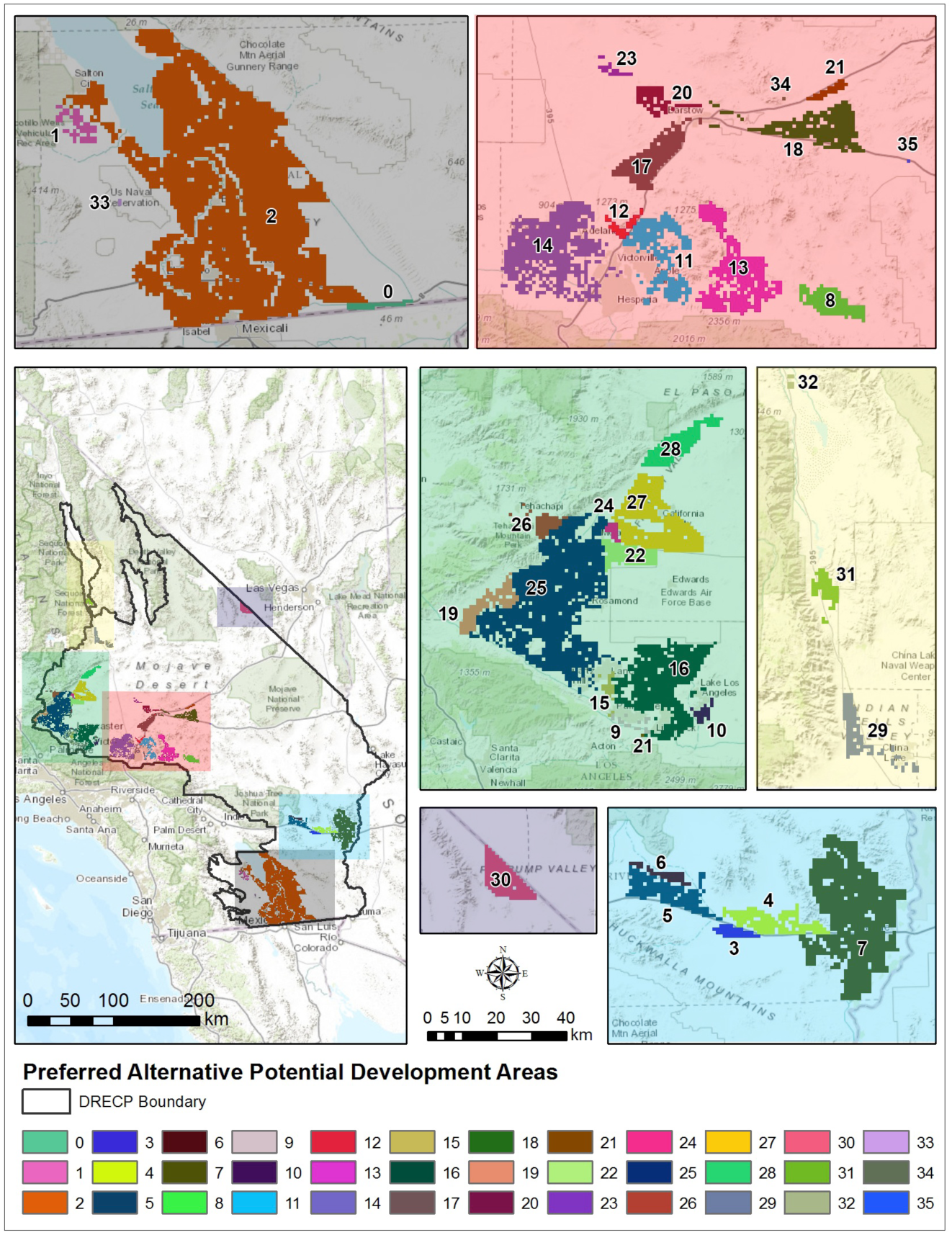
Spatial configuration of the proposed development chunks (see Appendix 4).

### Comparison of all chunks across all alternatives

Similarly, we compiled a ranking of proposed development chunks across all five alternatives. This information may be useful if the final development plan mixes and matches chunks from different alternatives. However, it is critical to note that the influence of developing certain regions of the landscape is not additive. Interactions between chunks, such as several chunks directly adjacent to one another that form a long barrier to gene flow, may have a significantly higher impact on gene flow than the sum of those individual chunks alone. The best available methodology for assessing a landscape-level development plan is to evaluate the effects of removing all of the proposed development chunks simultaneously. The results showing the individual impacts of the different putative development chunks across all of the alternatives can be found in Appendix 5.

## Discussion

The data we have generated for this project, including over 1.2 trillion base pairs of DNA sequence data and 80 high-resolution raster data layers covering much of the Mojave Desert will be useful for further work to understand the natural history and ecology of the desert tortoise and for guiding conservation actions.

There are several important caveats that should be kept in mind with respect to our approach. First, we model the development chunks, as we calculated them (see Appendix 4), as project areas with impenetrable boundaries that completely repel tortoise movement, but never kill or otherwise reduce tortoise fitness. Other modeling strategies are possible, and they will have different effects on the predictions of gene flow reduction or increase after development. Second, although we include the full geographical range of *G. agassizi* in our analysis, we only model development in the California portion of its range, and only that development identified in the possible DRECP scenarios. In particular, potential development in Nevada will have consequences for gene flow in California, and ideally scenarios for both states should be evaluated simultaneously to develop a comprehensive, range-wide picture of impacts on the species. Additionally, the impact of the reduction in gene flow that we model on ecological and demographic processes, and therefore on population viability, is not currently known. Modeling changes in population viability is a critical next step in our research.

Furthermore, although the methods we use represent the currently best available, the suitability of commute time as a proxy for divergence time (what we actually estimate from genome sequence) is currently unclear. Nonetheless, we do not think this discrepancy is responsible for the substantial difference between observed divergence times and those predicted under the model. The main unexplained discrepancy we observed was the north/south split; our best guess is that this is a result of historical barriers produced by habitat change and/or the existence of large, saline lakes in many of the basins in the area over the past tens of thousands of years. Further methods are required to incorporate a temporal dimension to the model we use. By fitting the model using only divergences between tortoises on the same side of the barrier, we hope to mostly exclude the effect of this deeper history.

Given these caveats, we feel that certain biological conclusions of key importance to Mojave desert tortoise conservation generally, and the DRECP in particular, can be made at this time.

1. By sampling the entire tortoise genome, we can detect subtle differences in population structure that have previously been impossible to detect with more conventional genetic and genomic tools (Figure 2), and is relatively free of statistical bias.
2. We detected a strong signal of isolation by distance among tortoises, and that signal is consistent across spatial scales and habitat regions across the range of the tortoise. Even within the relatively homogeneous Ivanpah Valley, we found a strong, statistically significant relationship between genetic and geographic distances. We conclude from this result that even tortoise populations within uninterrupted basins are not "panmictic", allowing the potential for local adaptation regionally in the desert. We also conclude that the occasional long-distance dispersal events that have been observed for the species do not seem to be leading to large areas of admixture and free interbreeding. Rather, there appears to be a general relationship between genetic isolation and geographic distance that scales across both large and small landscapes. The extent to which locally adapted populations are, or are not, exchangeable is an important avenue for future research, given the frequent use of translocations as a management tool in tortoises.
3. Our PCA analysis identified three primary groupings of tortoises, corresponding to a north/south division, and within the southern group, and east-west division (inset of Figure 4). Certain aspects of these groupings were also suggested in previous analyses (Hagerty and Tracy 2010), although the concordance is not perfect. In particular, the split between the California Cluster and the Las Vegas Cluster of Hagerty and Tracy is almost exactly replicated on PC1, and the NC, WM, and EC splits among the California Cluster are recovered by PC2. Our data indicate that the North-South groups comprise two key tortoise management units, and the East-West division is an additional genetic grouping. For the North-South grouping, our data suggest that the mountains separating the Ivanpah Valley from the surrounding desert habitat constitute a major barrier to gene flow. Within the southern unit, the Cadiz Valley (and its extension to the north and west, the Baker Sink, see Hagerty and Tracy 2010) has similarly been a low-elevation barrier to gene flow. Both of these suggest that if alternative energy installations could be placed *within* the New York/Providence mountains or the Cadiz Valley, they would interfere relatively little with current or past tortoise metapopulation dynamics, but that installations in the corridors of tortoise habitat around these could easily isolate these areas. Our data further indicate that these three sets of tortoises should best be considered three independent management units for tortoise conservation. Other units may emerge with additional tortoise sampling or analysis, but these three are clear.
4. Our landscape genetic inference framework allowed us to estimate the relative effects that the different development alternatives put forth in the DRECP will have on desert tortoise gene flow in the Mojave. Alternative 1 was found to have the least effect on tortoises, followed in order by the Preferred Alternative, Alternative 4, Alternative 3, and Alternative 2. However, we also note that the effects of all five of the alternative development plans have profound effects on tortoise gene flow; the additional time required for gene flow ranges from 650-950 years across alternative plans, and the relative isolation from 9% to 26%. These numbers far outstrip the actual amount of lost habitat for each plan (1.7%-4.9% of the total habitat for the tortoise), and emphasize the cascading effects that development can have on landscape connectivity.
5. Within the Preferred Alternative, many of the individual proposed development chunks have relatively little impact on desert tortoise connectivity when considered in isolation, although some have extremely high impacts. Chunks 31, 32, 25, and 7 have (in order) the greatest impact on tortoise connectivity, and should be examined closely before they are implemented. One of these, chunk 31 is located at the entrance of the Owens Valley, and this chunk is having a disproportionate effect on gene flow across the whole range of the tortoise, because it is effectively isolating the Owens Valley from the rest of the Mojave Desert. The general lesson from these analyses is that development that isolates regions should be avoided.
6. Across all alternatives, we identified and evaluated 214 development chunks, in terms of their individual effects on tortoise connectivity, and we encourage using this list along with other variables as a first pass for considering the order of approval of projects and habitat patches. However, we again emphasize that this list is for each chunk by itself, and it ignores the synergistic effects of developing multiple chunks.
7. Future analyses can, and should, investigate the isolating effects that multiple habitat chunks have when considered together. By sequentially adding development chunks and subdividing chunks, our work can help develop both a better final build-out and a gradual path to that build-out that minimized impacts on tortoise connectivity for as long as possible across the Mojave. Further direction from CDFW would be valuable in delineating which particular combinations of proposed development chunks would be useful to evaluate.
8. The tools we have developed can be used to predict the local effects of gene flow of specific development plans, and to recommend specific mitigation procedures, including critical issues like the location of habitat corridors. Not only selection of development areas (in the DRECP), but also situation of development within these areas in subsequent planning processes, will be key to reducing the impact of renewable energy development on the long-term viability of desert tortoise populations.

## Acknowledgements

We thank Roy Averill-Murray of the US Fish and Wildlife Service, Danna Hinderle, and the staff and scientists from the DTCC, and Jim Oosterhuis from the San Diego Zoo for desert tortoise handling training and strategizing on this project, Adalgisa Caccone, Kevin White, and Zifeng Jiang for sharing an early draft assembly of the Galapagos tortoise genome, Tasha La Doux, Jim Andre, and the Sweeney Granites Mountains Desert Research Center for hospitality and field support, Ying Zhi Lim, and Sarah Wenner for sorting tortoise samples, and Mariel Villanueva for some tortoise background research. This work was made possible through financial support from the California Department of Fish and Wildlife, Agreement Number P1382008. This work used the Vincent J. Coates Genomics Sequencing Laboratory at UC Berkeley, supported by NIH S10 Instrumentation Grants S10RR029668 and S10RR027303. PR and EL are supported by NSF grant #DBI-1262645 and startup funds from USC.

## Appendix 1 GIS Raster Data

We compiled and synthesized a total of 83 environmental raster data layers for this study area. These rasters fell in five principal categories: anthropogenic, biotic, climatic, topographic, and soils. We briefly describe these layers below. The resolution of the layers varied from 10m to 800m, although they were all standardized to 30m. For a list of all 83 layers, including a brief description of each, see Figure A7.

### Anthropogenic layers

Our anthropogenic GIS raster dataset included 13 layers, sourced or derived from the 2012 TIGER Census road classification (http://www.census.gov/cgi-bin/geo/shapefiles2012/main). The data describe the spatial distribution of roads in the Mojave ranging from 4WD trails and bike paths to primary roads and their on- and off-ramps. These layers were also used to calculate the Euclidean distance across the landscape to the nearest road. Also included was one layer from the National Land Cover Database 2011 (http://www.mrlc.gov/nlcd2011.php) that depicts the percent of impervious cover per cell. All anthropogenic layers have a resolution of 30m.

### Biotic layers

Our biotic GIS raster dataset included four layers describing the distribution of plant material across the Mojave. Three of these layers (shrub/scrub, grass/herb, and tree cover) were sourced from the National Land Cover Database 2011 (http://www.mrlc.gov/nlcd2011.php) and have a resolution of 30m. The fourth layer, annual growth potential, was calculated from the Moderate Resolution Imaging Spectroradiometer Enhanced Vegetation Index (MODIS-EVI) following the methods of Wallace and Thomas (2008) and Nussear *et al.* (2009). This 250m resolution layer serves as a proxy for annual plant biomass and was aggregated down to 30m resolution.

### Climatic layers

The climatic GIS raster dataset consisted of 52 layers describing the spatial distribution of climatic variables, including minimum, maximum, and mean temperature as well as mean precipitation, for each month and a yearly average, across the Mojave. These layers were taken from the PRISM Climate Group (http://www.prism.oregonstate.edu/normals/) and calculated over the last 30 years. The resolution of these layers was 800m, and were aggregated down to 30m.

### Topographic layers

The topographic GIS raster dataset consisted of 11 layers derived from the National Land Cover Database (NLCD; http://www.mrlc.gov/nlcd2011.php) and National Elevation Database (NED) on the USGS National Map Viewer (http://viewer.nationalmap.gov/viewer/). The elevation, aspect, surface roughness, surface area, slope, and eastness and northness (the degree to which slope faces east and north, respectively) layers were all derived from the NED DEM. The land cover and barrenness layers were both extracted from the NLCD. All landscape raster layers were produced at a 30m resolution. Longitude and latitude rasters were also constructed over the study area.

### Soil layers

The soils GIS raster dataset consisted of 3 layers that describe bulk density, percent of rocks greater than 10 inches, and depth to bedrock. The data were extracted from SSURGO2 database. Data gaps in SSURGO2 were filled using STATSGO, downloaded from USDA NRCS Soil Data Mart (http://websoilsurvey.sc.egov.usda.gov/App/WebSoilSurvey.aspx). All layers were transformed to 30m rasters.

Of this complete set of rasters, three different subsets (of 6, 12, and 24 rasters) were selected to be used in inference of the correlation between ecological heterogeneity and partitioning of genetic variation in Mojave tortoises. We selected these subsets to reduce the complexity and computation time of analyses, and to produce a set of statistically more independent layers. Many of the initial 83 raster layers in the full set were highly correlated (Figure A8), and including such highly correlated layers in later analyses can both confuse the analysis and make any interpretation of their individual effects difficult or impossible. We selected rasters for inclusion to maximize overlap with layers used in previous GIS analyses of tortoises, and to minimize pairwise correlation among layers. The list of rasters included in each subset can be found in Figure A9.

### Appendix 2

#### Divergence

Suppose that we have sequences of length *L* from two individuals, with *C*_*i,k*_ reads that map for individual *k* to a position overlapping site *i*, for *k* ∈ {1, 2} and 1 ≤ *i* ≤ *L*. Suppose that *C*_*i,k*_ is marginally Poisson with mean *λ*_*k*_*c*_*i*_, and that each read covering site *i* independently draws an allele, so that given *C*_*i,k*_, the allele counts *N*_*i,k*_(*a*) are Multinomial with probabilities *p*_*i,k*_(*a*), where *a* is the allele. Suppose also that coverages and counts are independent between samples, given *p* and *c*. The probability that a pair of reads drawn from those at site *i*, one drawn uniformly at random from each sample, both have allele *a* is *Y*_*i*_(*a*)= *N*_*i,*1_(*a*)*N*_*i,*2_(*a*)*/C*_*i,*1_*C*_*i,*2_, and so that if *C*_*i,*1_*C*_*i,*2_ *>* 0,

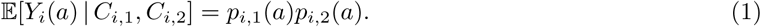

We would like to estimate mean sequence divergence,

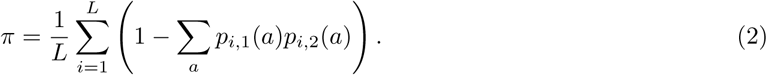

Given a weighting function *w* with *w*(0*, n*)= *w*(*n,* 0) = 0 for each *n*, an estimator of divergence is

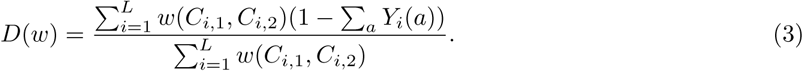

The expection of *D*(*w*) is

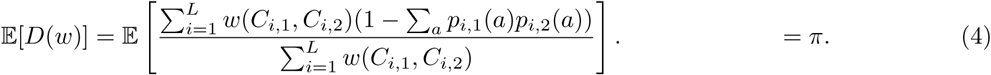

If the mean sitewise coverages *c*_*i*_ are independent of the *p*_*i,k*_, then by exchangeability, 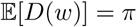. We take *w*(*x, y*)= *xy*:, which approximately (but not quite) does not depend on the coverages *λ*:

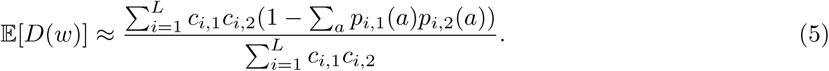

### Appendix 3

#### Model Specification

A model of landscape resistance, as discussed above, is essentially a specification of a reversible random walk on the landscape. A reversible random walk is specified by two quantities: the *stationary distribution* of each point *x*, denoted *π* (*x*), and the *relative jump rates* between each pair of adjacent locations *x* and *y*, denoted *j* (*x*, *y*); these combine to give the total rate of movement from *x* to *y* as *G* (*x*, *y*) = *j* (*x*, *y*) / *π* (*x*). The requirement that the random walk to be reversible, i.e. *π* (*x*) *G* (*x*, *y*) = *π* (*y*) *G*(*x*, *y*), means that relative jump rates must be symmetric, i.e. *j* (*x*, *y*) = *j* (*y*, *x*).

We then allow these two ingredients to be determined by linear functions of the landscape layers: if we have *n* landscape layers whose values at location *x* are *L*_1_ (*x*), …, *L*_*n*_ (*x*), then we suppose that

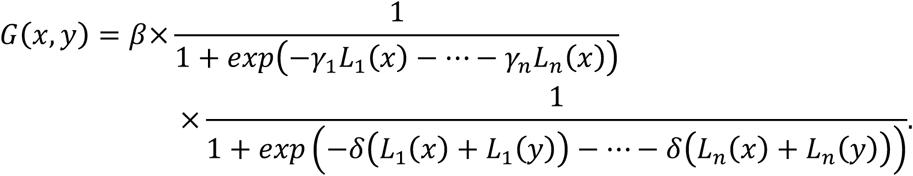

The parameters are: *β*, an overall scaling factor, and for each 1 ≤ *k* ≤ *n*, *γ*_*k*_, that determines how the *k*th layer affects the stationary distribution, and *δ*_*k*_, that determines how the *k*th layer affects the relative jump rates.

In practice, then, a model is determined by:

1. A *mask*, i.e. a specification of the total potential habitat area available for movement; movement rates to locations outside of this are assumed to be zero.
2. The *layers*, which provide a numerical value for each location on the landscape; we include a "constant" layer (that takes the value 1 everywhere), and normalize remaining layers to have mean zero and variance 1.
3. The *parameters β*, *γ*_1_, …, *γ*_*n*_, and *δ*_1_, …, *δ*_*n*_.
4. A *neighborhood size R* and a *local coalescence time T*.

These are combined to fit the data by computing for each *x* and *y* the mean time until a random walk begun at *x* first gets closer than *R* to the location *y*, which we denote by *h*_*R*_ (*x*, *y*), and postulating that the observed sequence divergence between tortoises at locations *x* and *y*, denoted *d* (*x*, *y*), is equal to *T* plus the mean *R*-commute time, i.e.

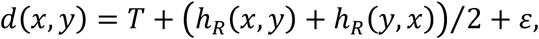

where *ε* is the noise due to demographic stochasticity and sequencing error.

#### Fitting Procedure

To fit the model above, we find parameters to minimize the weighted mean squared error

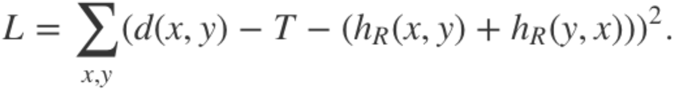

This requires computing the times *h*_*R*_ (*x*, *y*), which can be done as follows. First, we compute the movement rates of the random walk and place them in a matrix *G*, with rows and columns indexed by locations, and whose (*x*, *y*)th entry is *G*_*x*,*y*_ defined above. Fix a location *y* and a distance *R*, let *N*_*R*_ (*y*) be the set of locations within distance *R* of location *y*. Then the times *h*_*R*_ (*x*, *y*) solve the equations

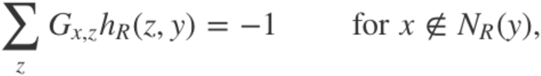

and boundary conditions

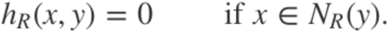

This forms one system of equations for each *y*, that we solve numerically using sparse matrix solvers in the *Matrix* package in *R* (Bates and Maechler 2014).

Analytically, the solution can be written as follows: for a given *y* and *R* let 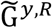 y,R denote the matrix obtained by removing the rows and columns of *G* corresponding to *N*_*R*_ (*y*). Then, seen as a vector indexed by *x*,

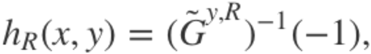

where 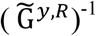 is the matrix inverse of 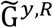 and (‒1) denotes the vector whose entries are all -1. This can be substituted into the expression for the mean squared error above, and then differentiated, to find analytic expressions for the gradient vector and Hessian matrix of *L* with respect to *T*, *β*, each *γ*, and each *δ*. With these in hand, we then use a "trust region" optimization routine, as coded in the package *trust* in *R* [Geyer]. This allows us to find best-fitting choices of all parameters except *R*; in practice, we then fix *R* at 15km. It would be preferrable to also optimize over *R*; however, *R* is nearly confounded with *T*, in that increasing *R* is very nearly equivalent to adding a constant to *h*_*R*_ (*x*, *y*), and so this choice does not significantly affect results.

#### Landscape Resistance Models

Concretely: for the *i*th sampled tortoise, let *n*_*i*_ be the number of other sampled tortoises within 25km, and let the (*i*, *j*)th *weight* be

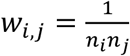, if *i ^ j* are ∈ the same region, and *w*_*i*,*j*_ = 0 otherwise. Let the (*i*, *j*)th *residual* be

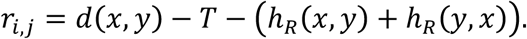

Then the *weighted median residual* is the value 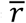 such that the sum of the weights of the residuals smaller then 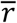 is equal to the sum of the weights larger than 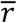: concretely, it satisfies

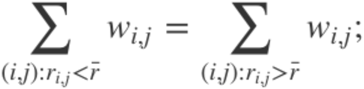

if there is ambiguity in where 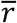 should fall, then it is specified as the weighted mean of the nearest possible samples.

### Appendix 4

#### Aggregation of development focus area polygons into “chunks”

The potential development areas evaluated in this study were derived from the alternative shapefiles on the DRECP gateway. Polygons designated as "Development Focus Areas", which ranged between 1000-2500 polygons for the different alternatives. The polygons were grouped into “chunks” according to area and proximity amongst each other, as described below.

First, an initial set of development chunks was established by selecting all polygons with an area greater than or equal to 1000 hectares. Next, all remaining polygons within 5 km (edge to edge proximity) of an initial polygon were identified as secondary polygons and assigned to the closest initial polygon. If the secondary polygons were adjacent to multiple initial polygons, the secondary polygon took the assignment of the largest initial polygon.

The remaining polygons (polygons under 1000 hectares that are not within 5 km of an initial polygon) were grouped into “remainder clusters,” based on proximity, with a 5 km upper limit. If a remainder cluster was smaller than 5 hectares and within 10 km of other such polygon clusters, that remainder cluster was reassigned to reflect a single potential development area with the closest remainder cluster. In the preferred alternative, the polygons were grouped into 36 potential development areas, with areas ranging between 4 and 300,000 hectares. All results were rasterized to a 1 km resolution (consistent with the habitat model from Nussear *et al.* (2009)).

### Appendix 5

#### Development chunks across all alternatives, ranked by isolation (years)

Below is a table of the effects of removing each chunk *across all alternatives*. Keys that show the spatial configuration all of the chunks can be found in Figures 10 and A8-A11). The *mean isolation* is a good measure of the total effect of removing a given area, but note that the absolute size of the number is not necessarily reflective of the overall effect, as it measures this piece, in isolation, without the effects of all other pieces. There are many instances where the impacts of multiple chunks considered together have a larger impact than the sum of the chunks by themselves.

- *habitat removed* is the total amount of area either in possible development areas or completely isolated from the rest of tortoise habitat under this alternative,
- *isolated* is the total area on which the gene flow to nearby areas has increased,
- *isolation* is the mean amount by which the commute time has increased to nearby areas across this area where it has increased,
- *max isolation* is the maximum amount by which the commute time has increased between any two nearby reference locations.

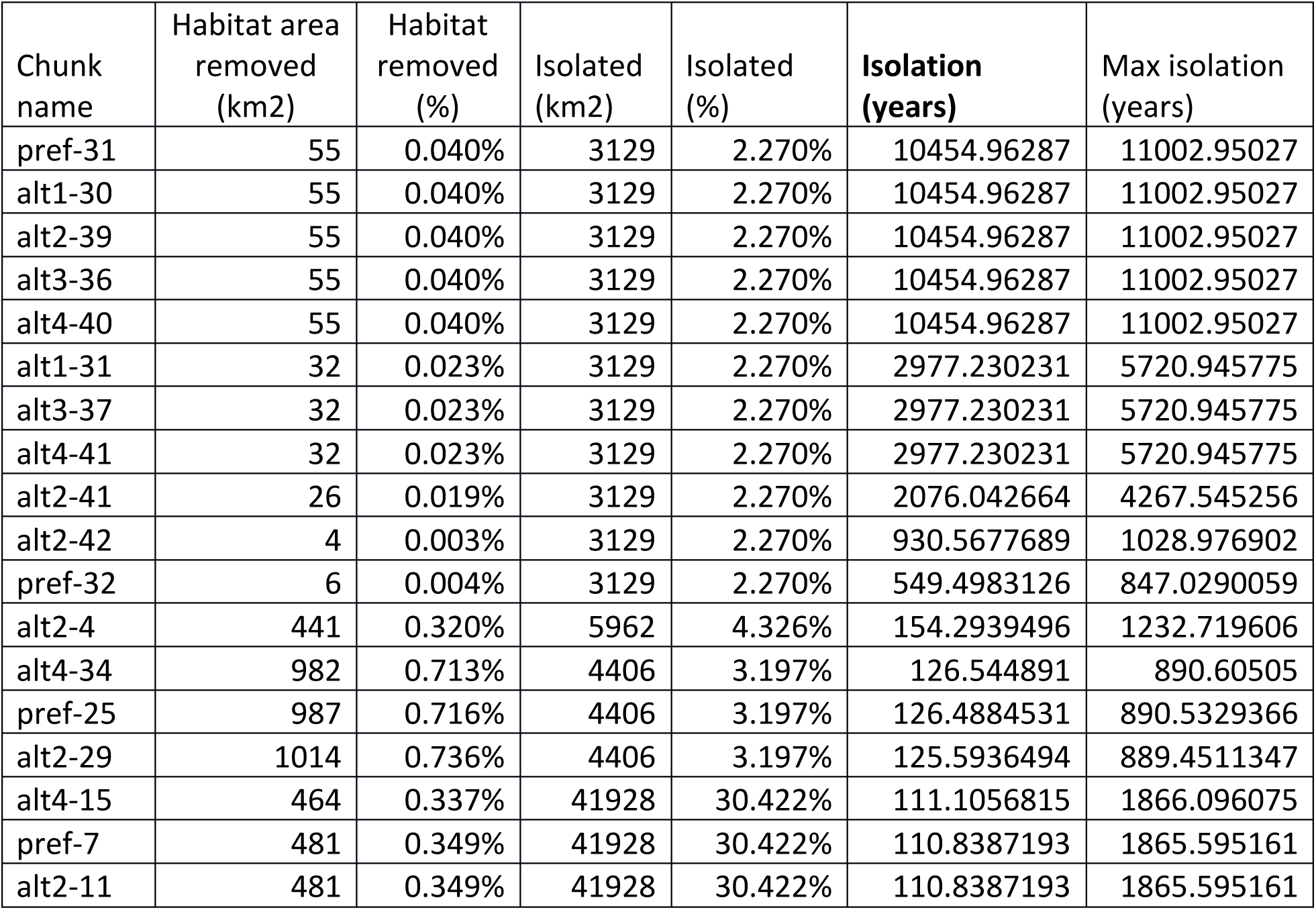

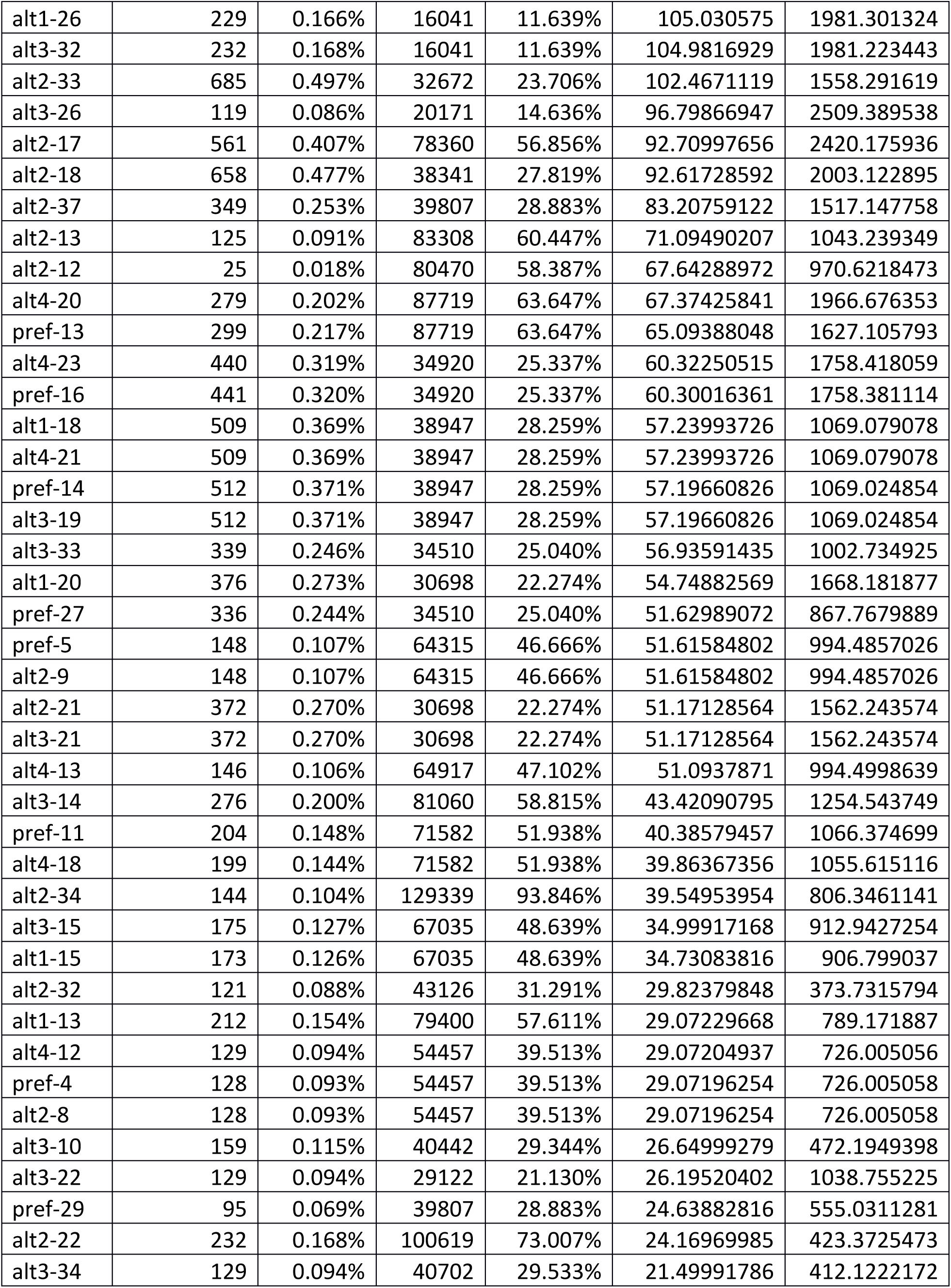

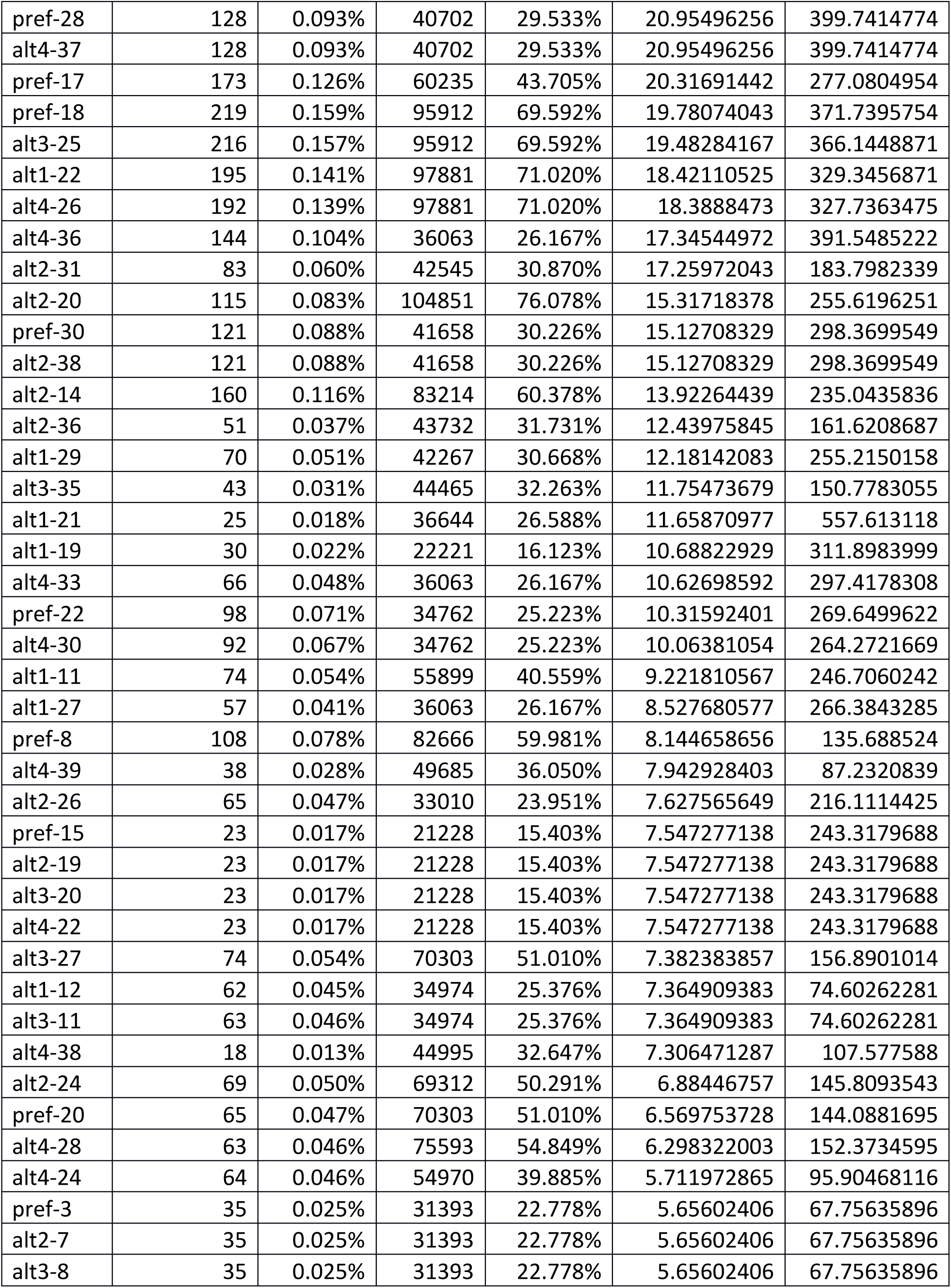

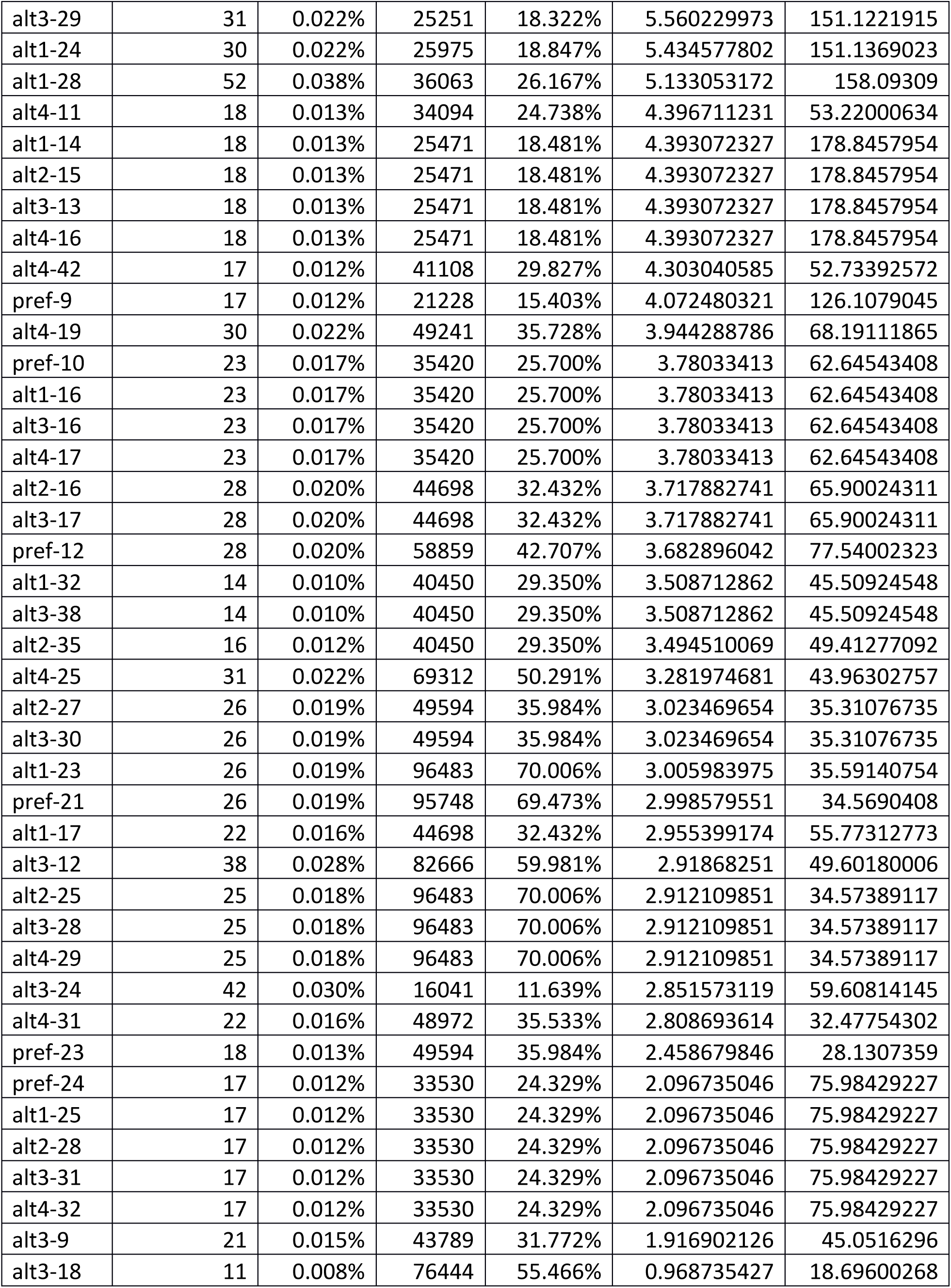

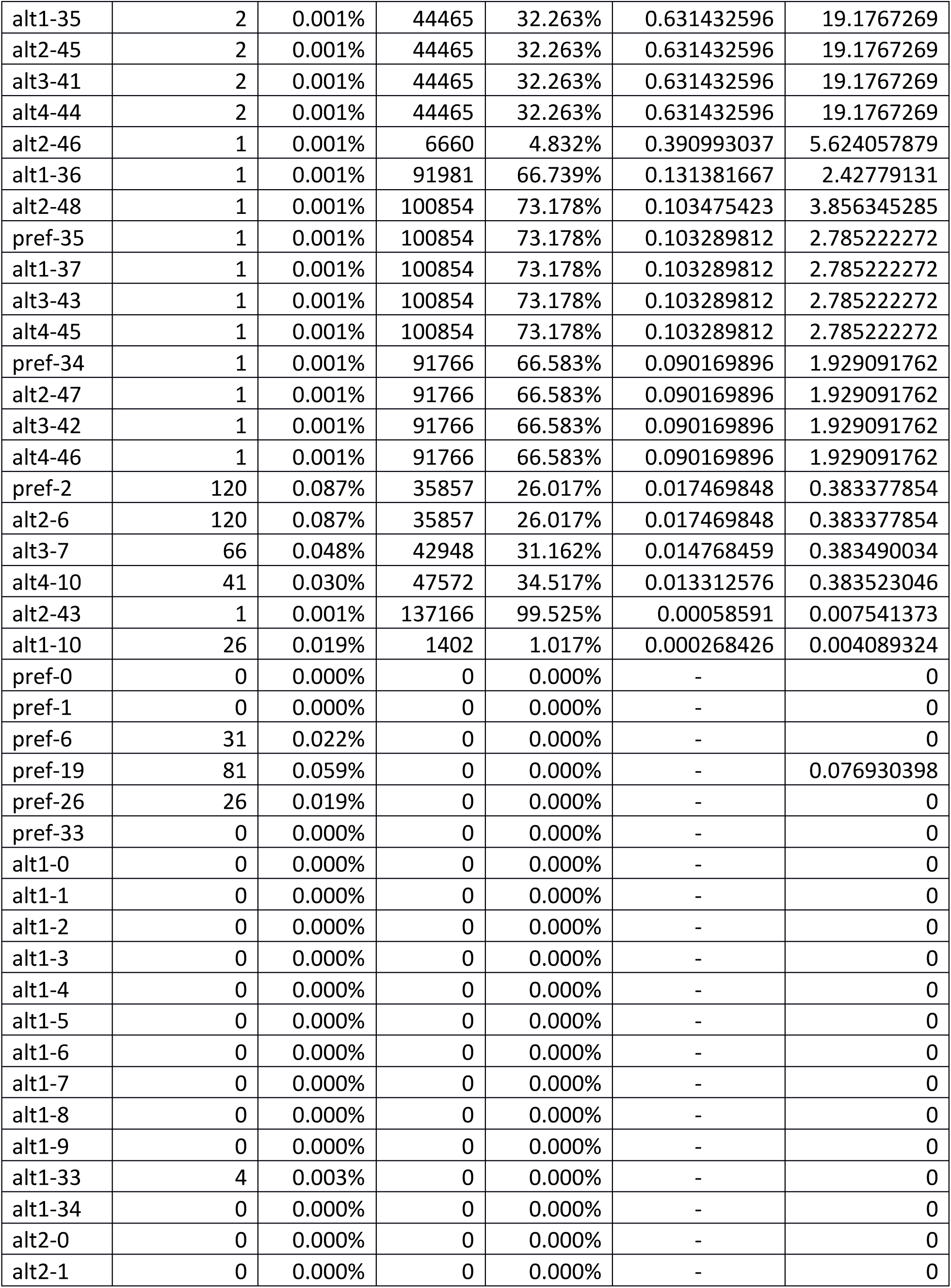

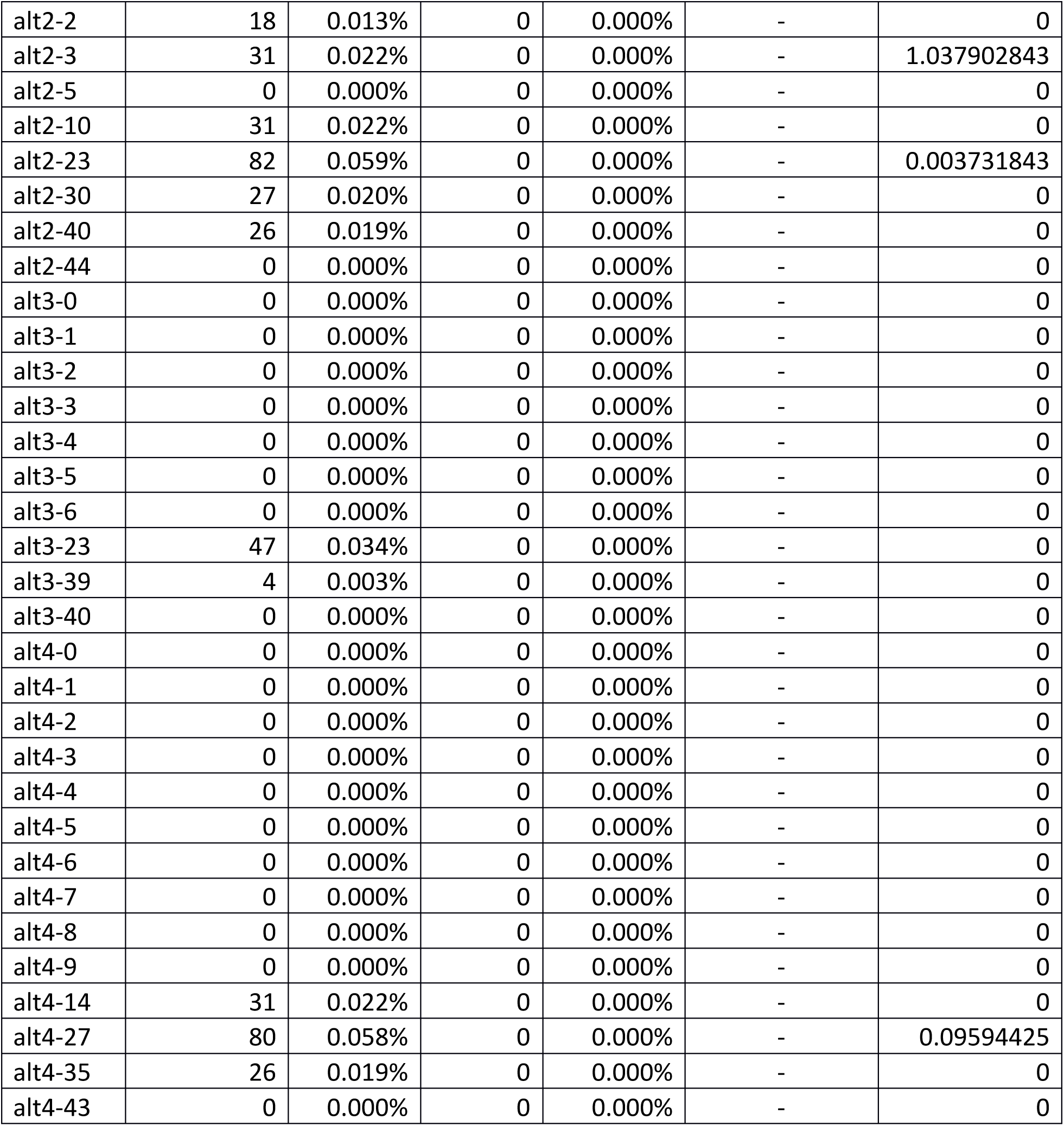

### Supplemental Figures

**Figure A1.**
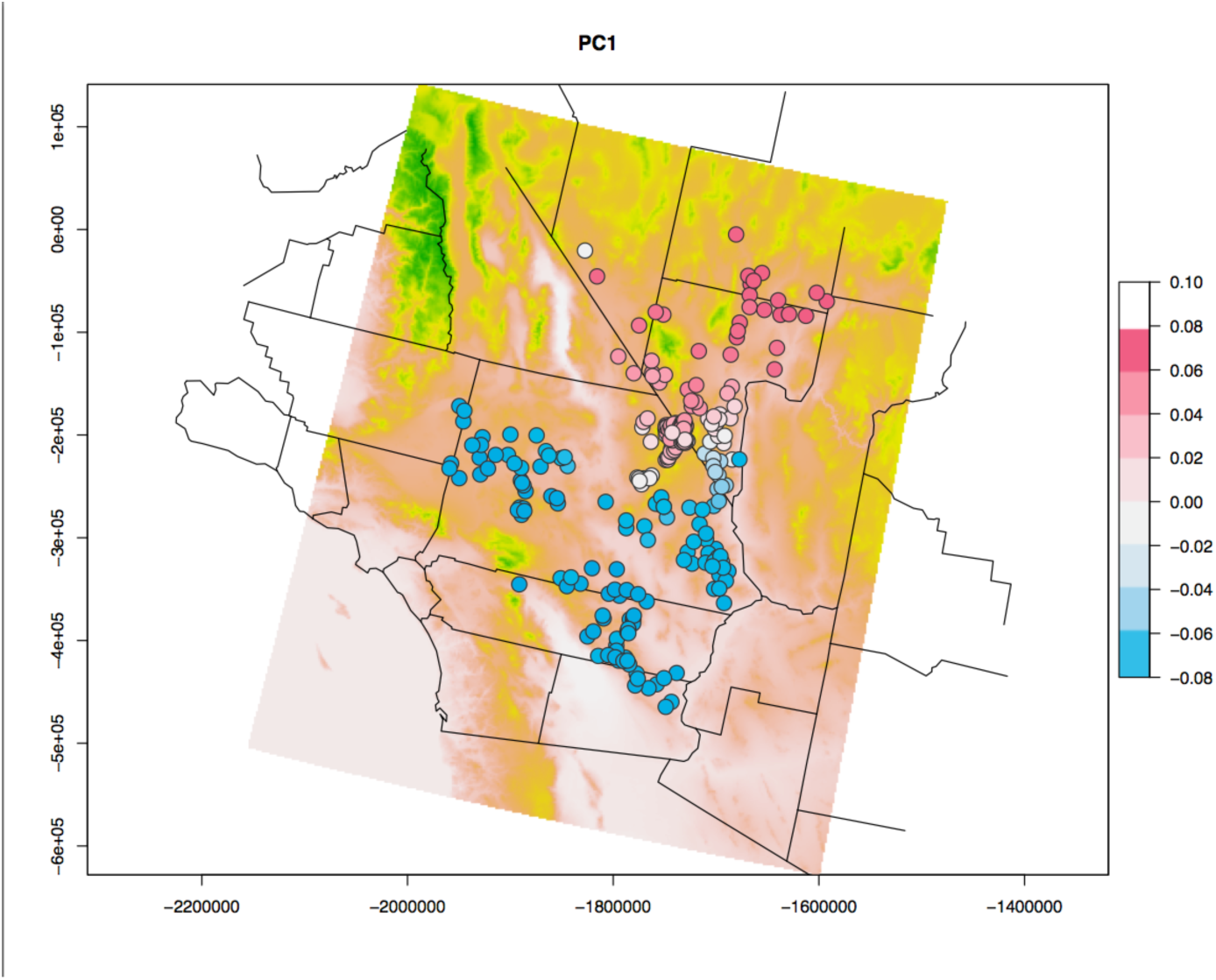
Visualization of genetic structure. Dots represent individual tortoises, and they are colored by their scores on PC1. Background color corresponds to elevation.

**Figure A2.**
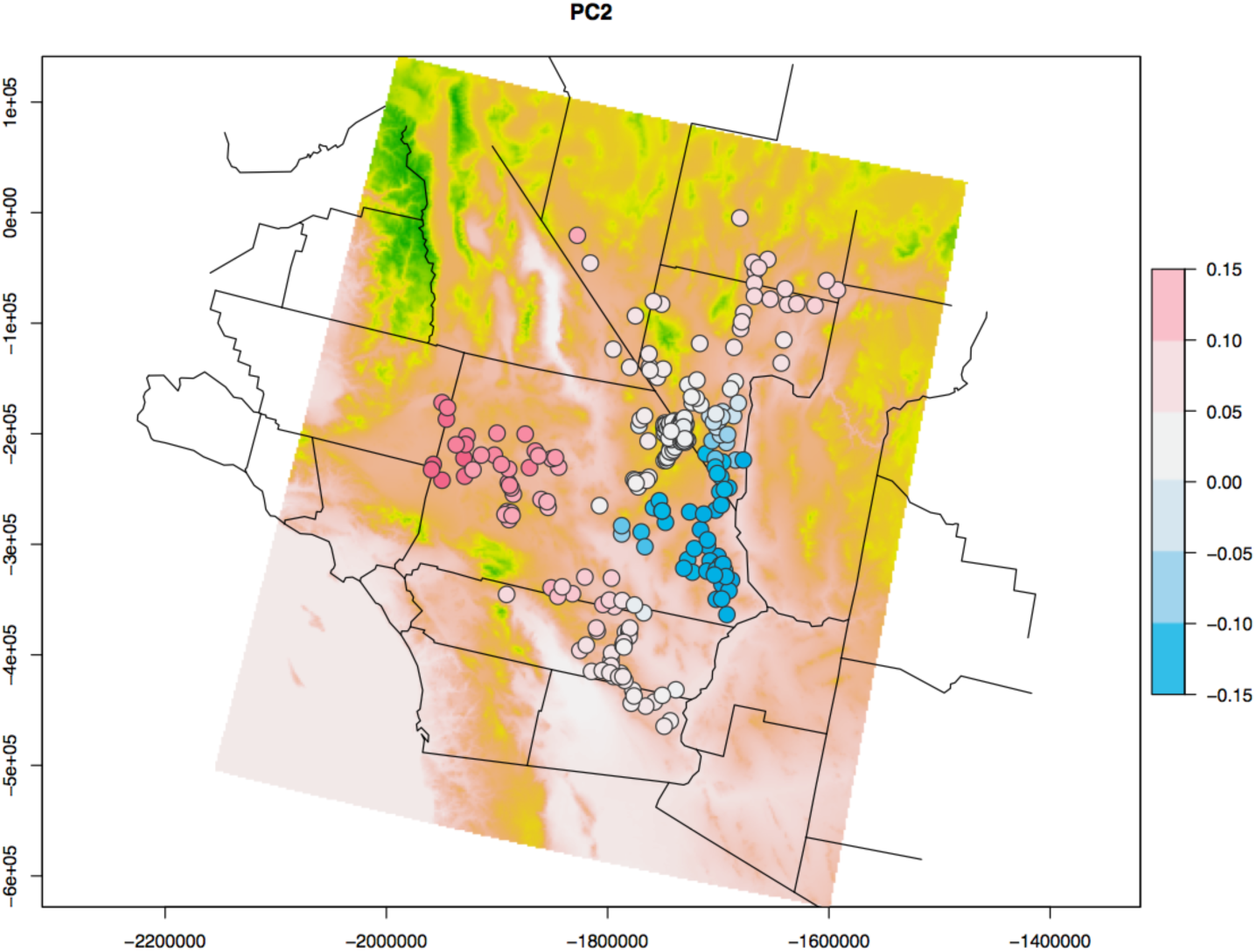
Visualization of genetic structure. Dots represent individual tortoises, and they are colored by their scores on PC2. Background color corresponds to elevation.

**Figure A3.**
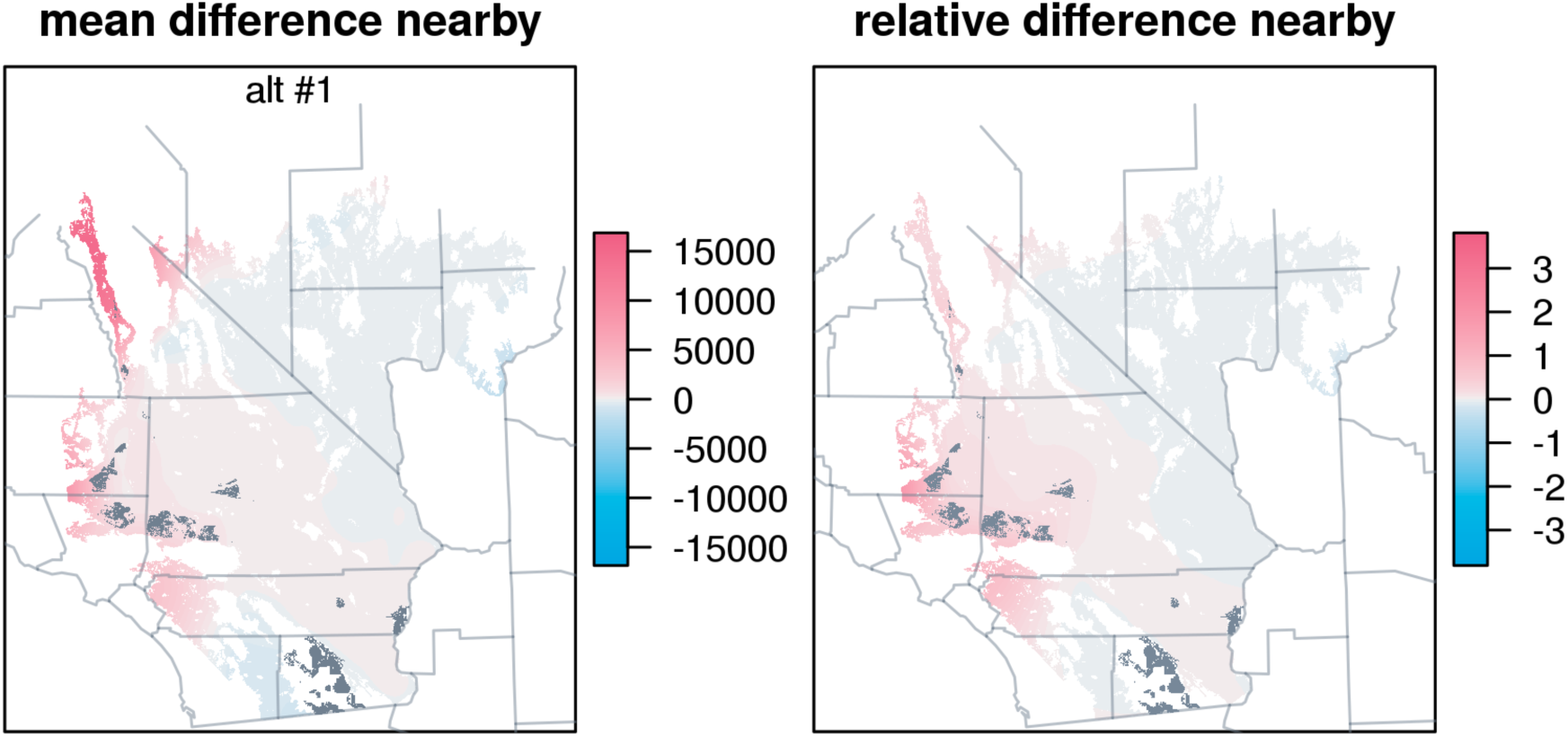
Effects of removing the development areas in Alternative 1.

**Figure A4.**
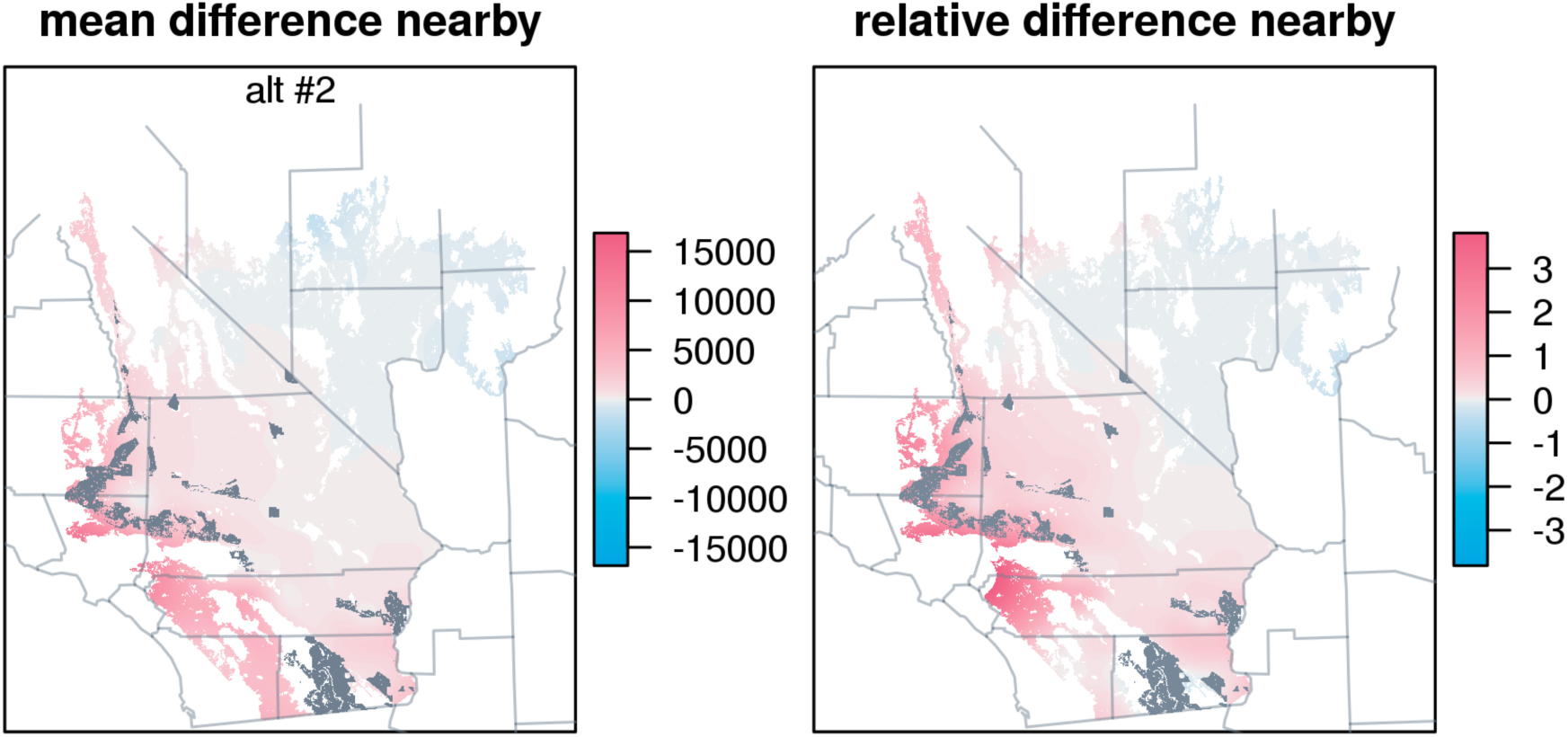
Effects of removing the development areas in Alternative 2.

**Figure A5.**
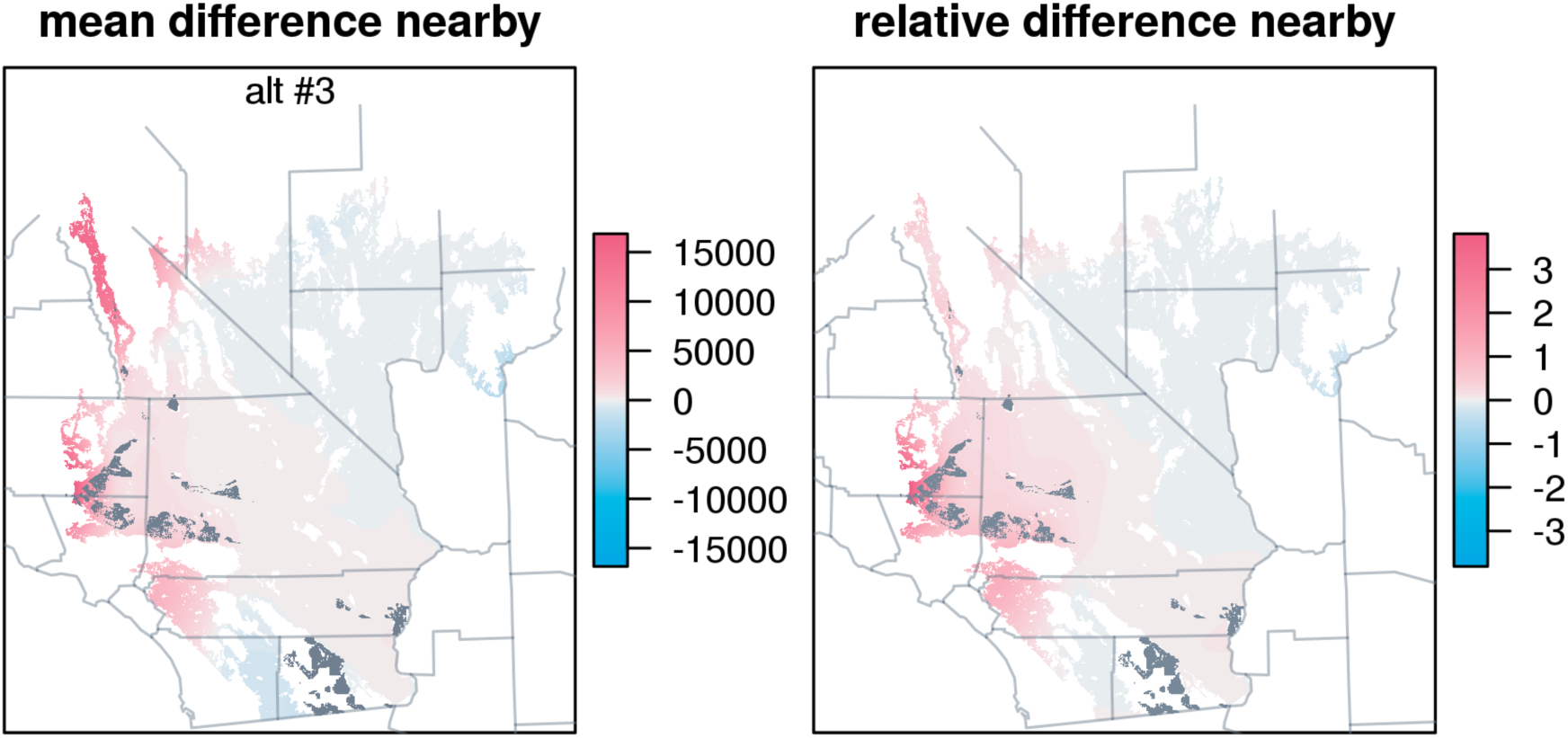
Effects of removing the development areas in Alternative 3.

**Figure A6.**
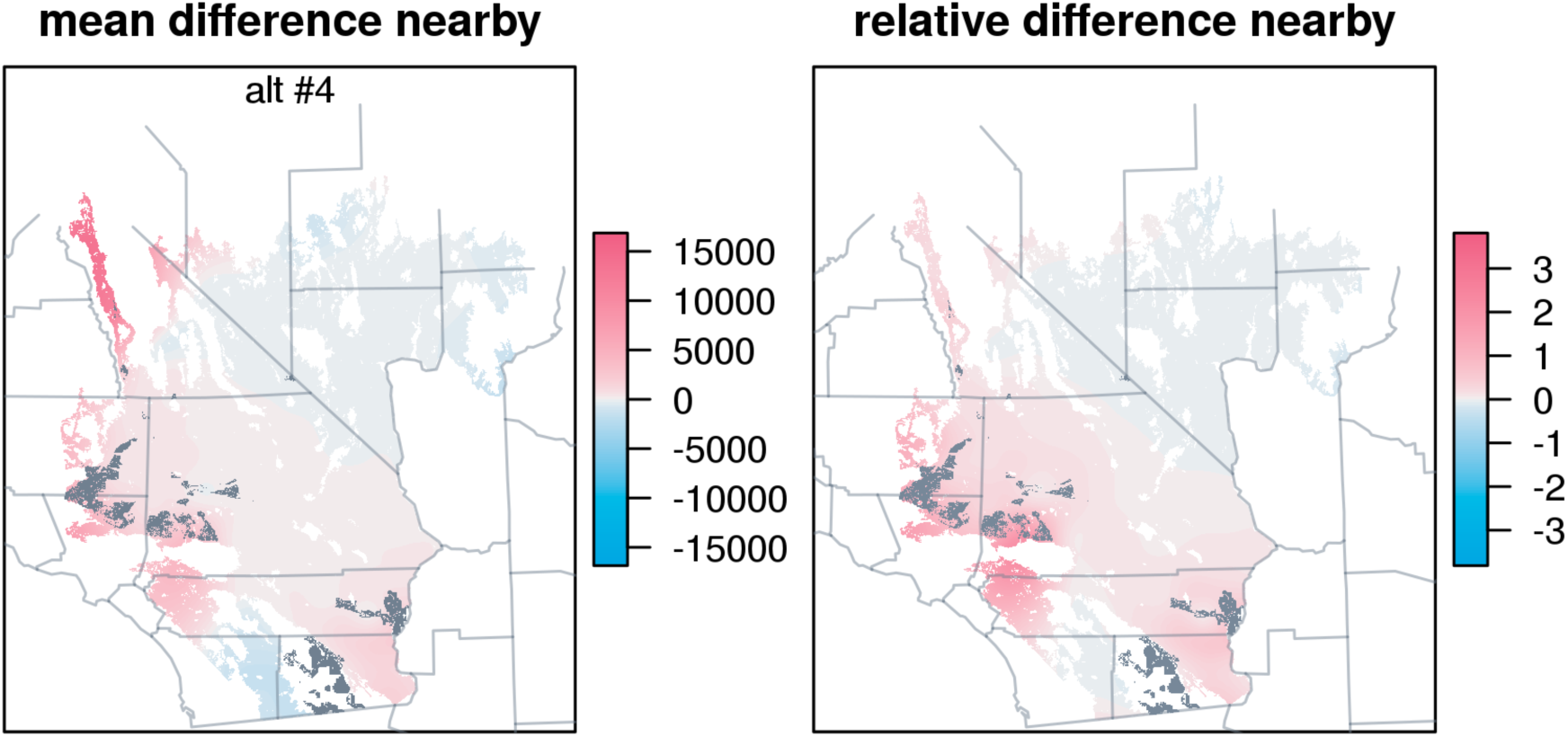
Effects of removing the development areas in Alternative 4.

**Figure A7.**
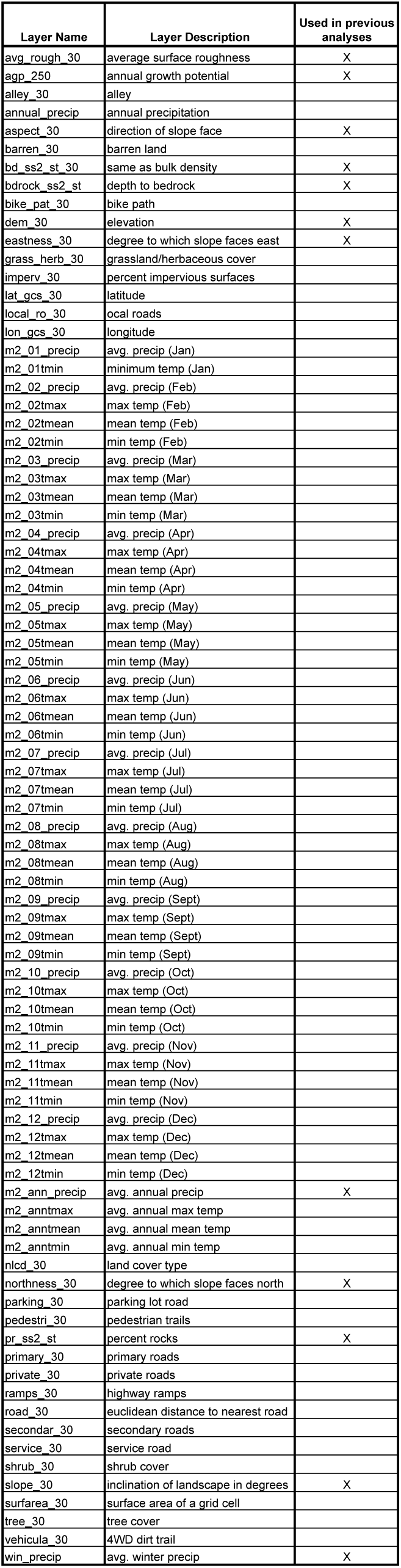
List of 83 landscape layers.

**Figure A8.**
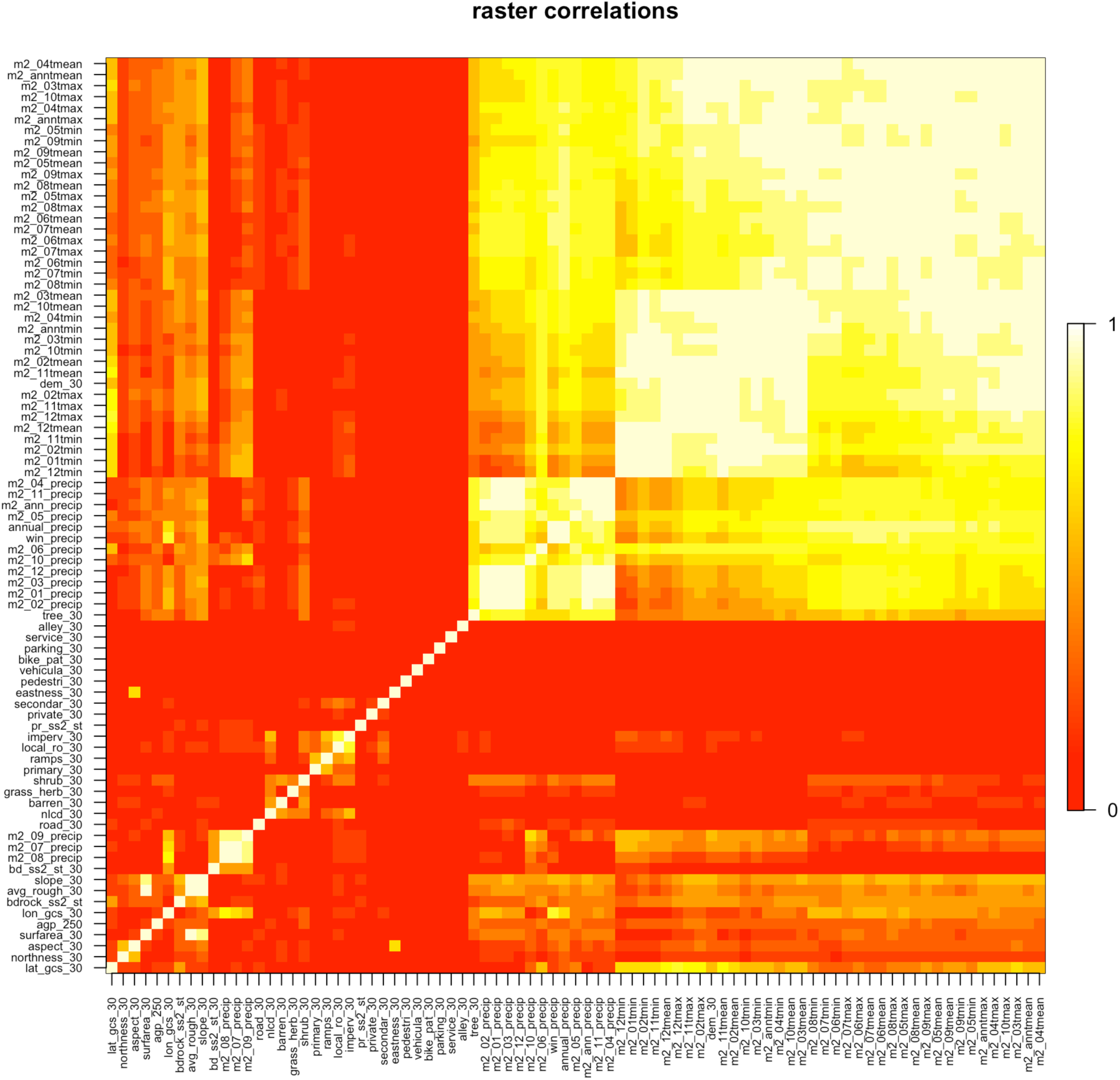
Matrix of correlations between spatial data layers.

**Figure A9.**
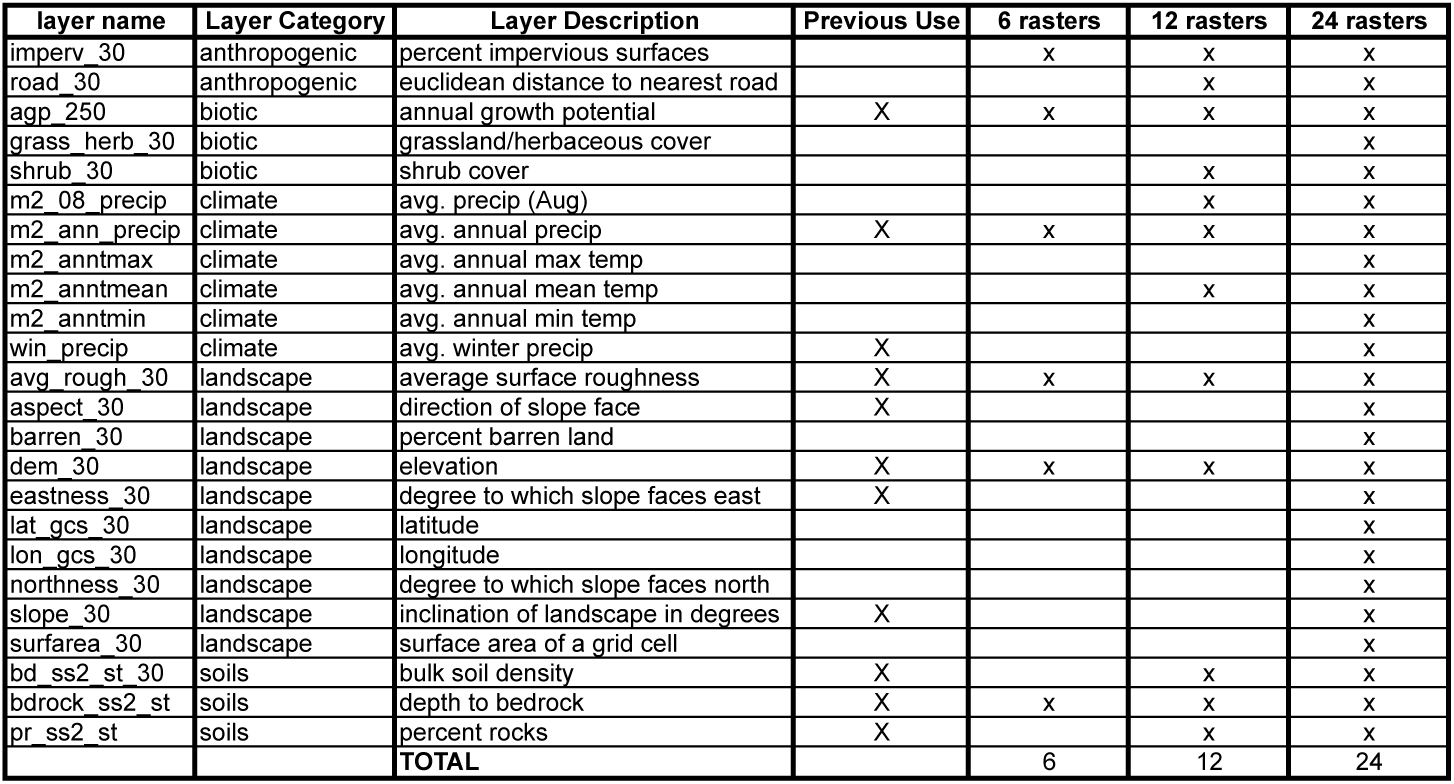
List of environmental data layers evaluated in 6, 12, and 24-layer models.

**Figure A10.**
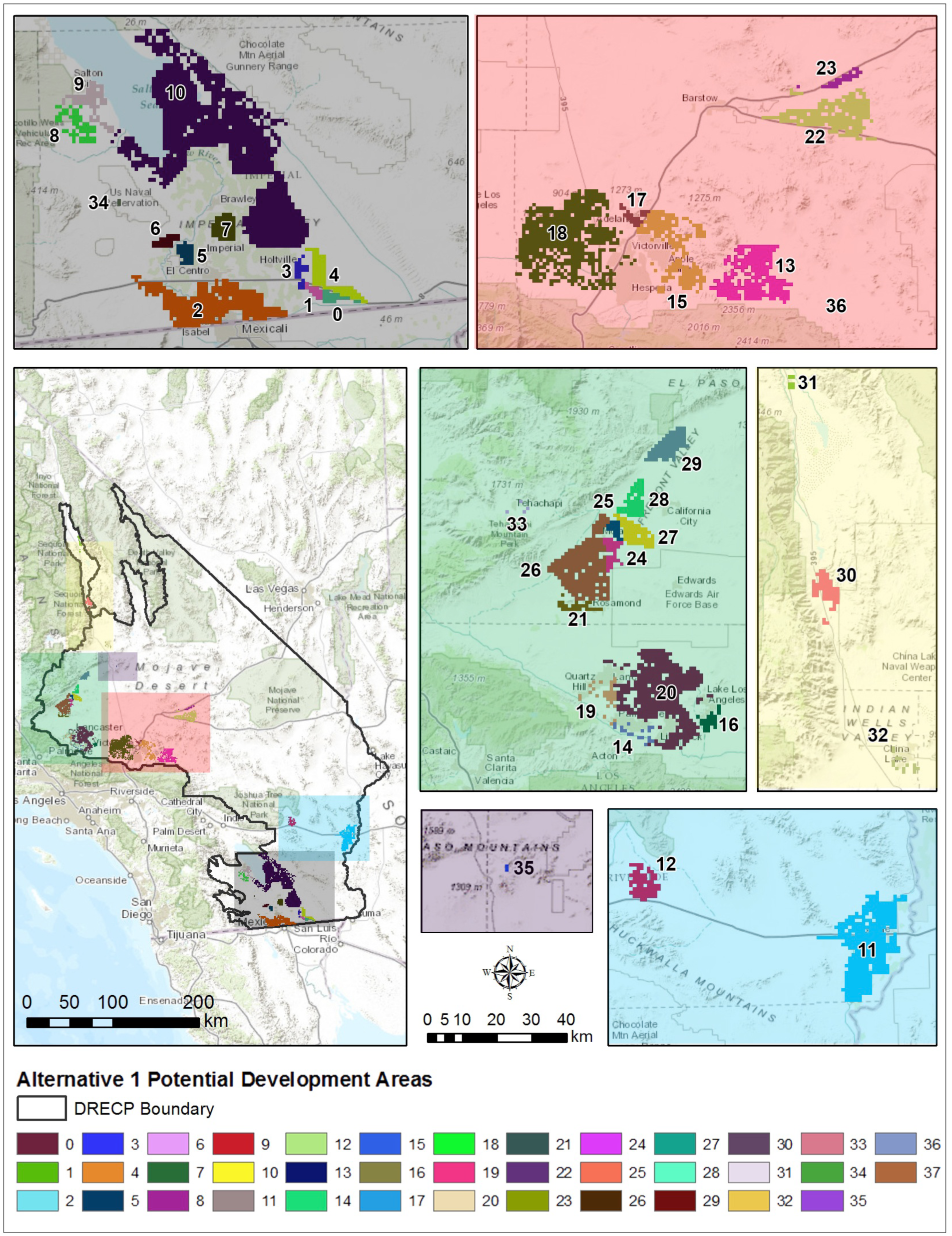
Spatial configuration of proposed development chunks in Alternative 1 that we analyzed.

**Figure A11.**
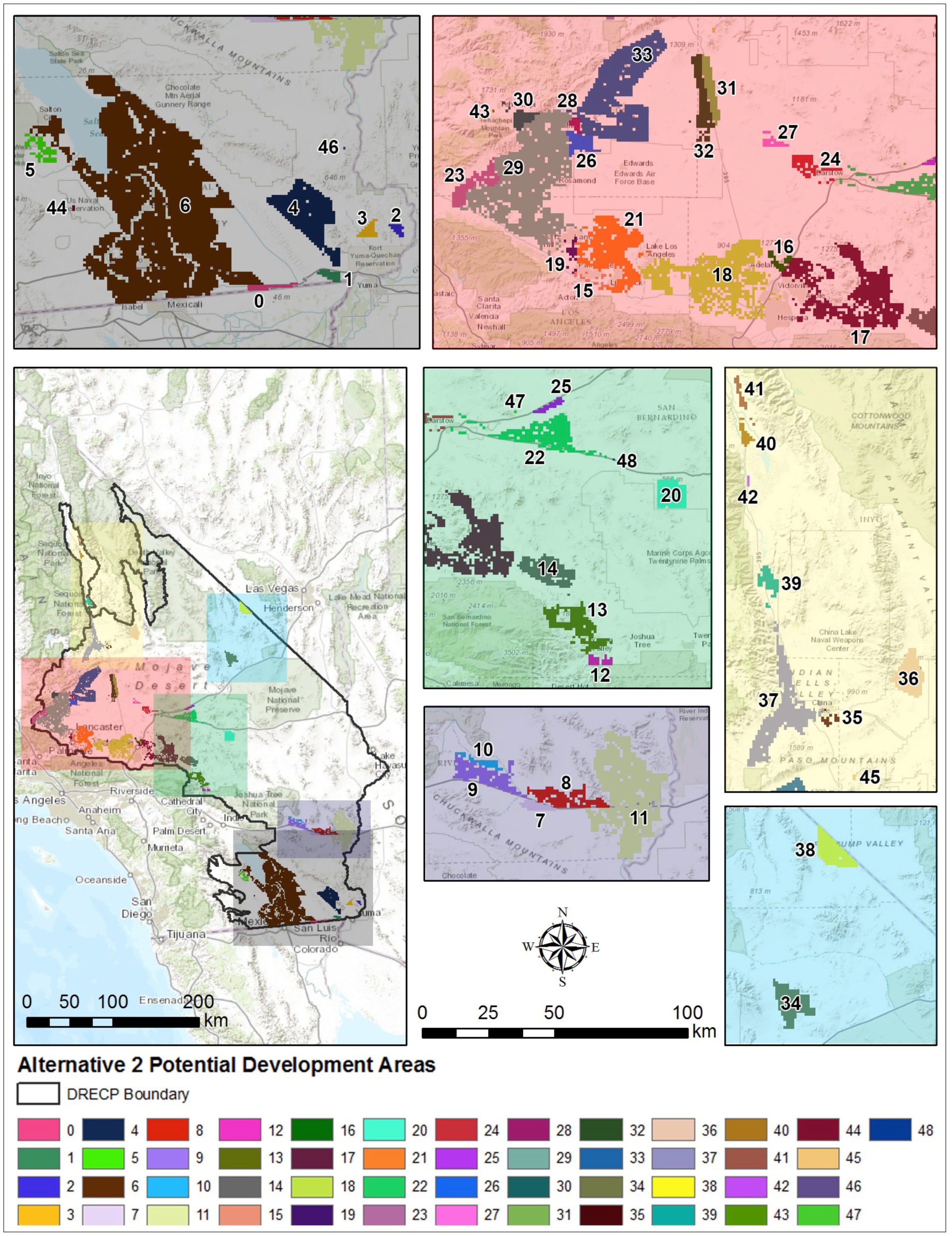
Spatial configuration of proposed development chunks in Alternative 2 that we analyzed.

**Figure A12.**
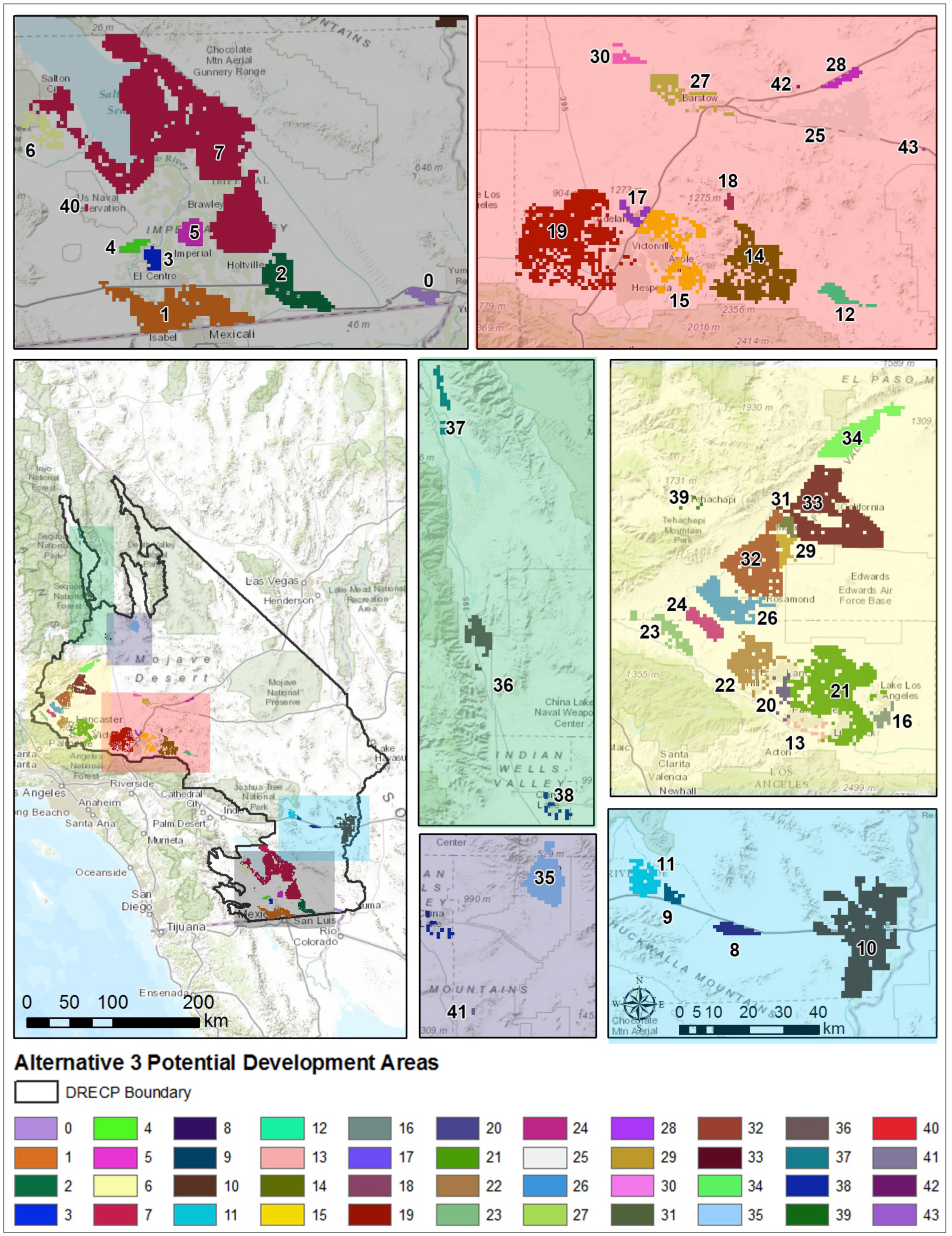
Spatial configuration of proposed development chunks in Alternative 3 that we analyzed.

**Figure A13.**
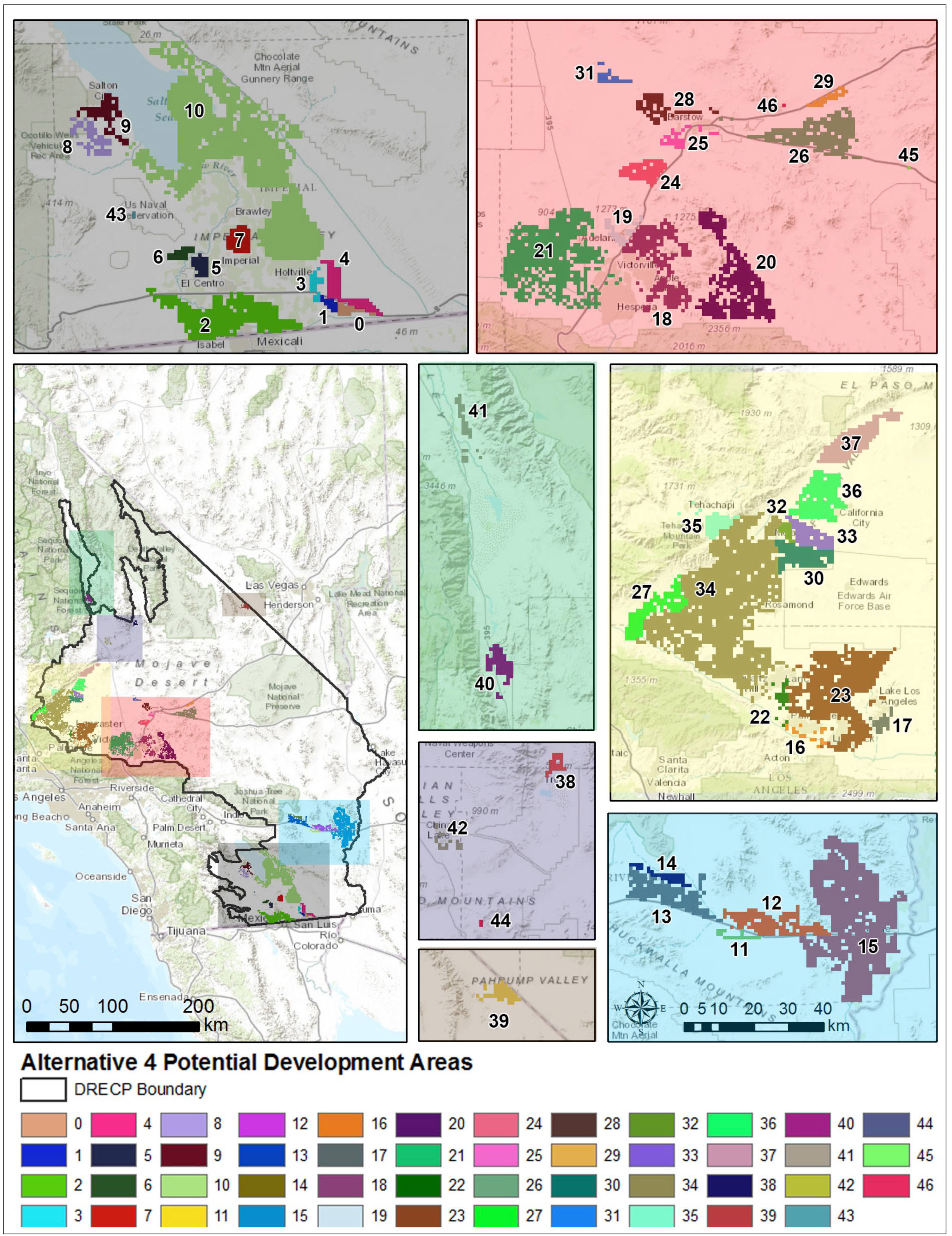
Spatial configuration of proposed development chunks in Alternative 4 that we analyzed.

